# Quantitative glycoproteomics reveals cellular substrate selectivity of the ER protein quality control sensors UGGT1 and UGGT2

**DOI:** 10.1101/2020.10.15.340927

**Authors:** Benjamin M. Adams, Nathan P. Canniff, Kevin P. Guay, Ida Signe Bohse Larsen, Daniel N. Hebert

## Abstract

UDP-glucose: glycoprotein glucosyltransferase (UGGT) 1 and 2 are central hubs in the chaperone network of the endoplasmic reticulum (ER), acting as gatekeepers to the early secretory pathway yet little is known about their cellular clients. These two quality control sensors control lectin chaperone binding and glycoprotein egress from ER. A quantitative glycoproteomics strategy was deployed to identify cellular substrates of the UGGTs at endogenous levels in CRISPR-edited HEK293 cells. The seventy-one UGGT substrates identified were mainly large multidomain and heavily glycosylated proteins when compared to the general N-glycome. UGGT1 was the dominant glucosyltransferase with a preference towards large plasma membrane proteins whereas UGGT2 favored the modification of smaller, soluble lysosomal proteins. This study sheds light on differential specificities and roles of UGGT1 and UGGT2 and provides insight into the cellular reliance on carbohydrate-dependent chaperone intervention by UGGT1 and UGGT2 to facilitate proper folding and maturation of the cellular N-glycome.

## INTRODUCTION

Protein folding in the cell is an error-prone process and protein misfolding is the basis for a large number of disease states (Hebert and Molinari, 2007; Hartl, 2017). A significant fraction of the proteome in mammalian cells passes through the secretory pathway by first being targeted to the endoplasmic reticulum (ER) where folding occurs (Uhlén et al., 2015; Itzhak et al., 2016; Adams et al., 2019a). Molecular chaperones of the ER help to guide secretory pathway cargo along a productive folding pathway by directing the trajectory of the folding reaction, inhibiting non-productive side reactions such as aggregation or by retaining immature or misfolded proteins in the ER until they can properly fold or be targeted for degradation. Understanding how chaperone binding controls the maturation and flux of proteins through the secretory pathway is of important fundamental biological concern and will impact our knowledge of protein folding diseases and the development of potential therapeutics including the production of biologics that are frequently secretory proteins.

Proteins that traverse the secretory pathway are commonly modified with N-linked glycans as they enter the ER lumen (Zielinska et al., 2010). These carbohydrates serve a variety of roles including acting as quality control tags or attachment sites for the lectin ER chaperones calnexin and calreticulin (Helenius and Aebi, 2004; Hebert et al., 2014). N-glycosylation commences co-translationally in mammals and the first round of binding to calnexin and calreticulin is initiated shortly thereafter by the rapid trimming of glucoses by glucosidases I and II to reach their monoglucosylated state (Chen et al., 1995; Cherepanova et al., 2019). Lectin chaperone binding is multifunctional as it has been shown to: (1) direct the folding trajectory of a protein by acting as a holdase that slows folding in a region-specific manner; (2) act as an adapter or platform to recruit folding factors including oxidoreductases (ERp57 and ERp29) and a peptidyl-prolyl *cis trans* isomerase (CypB) to maturing nascent chains; (3) diminish aggregation; (4) retain immature, misfolded or unassembled proteins in the ER; and (5) target aberrant proteins for degradation (Rajagopalan et al., 1994; Hebert et al., 1996; Daniels et al., 2003; Molinari et al., 2003; Oda et al., 2003; Wang et al., 2008; Kozlov and Gehring, 2020). For glycoproteins, the lectin chaperones appear to be the dominant chaperone system as once an N-glycan is added to a region on a protein, it has been shown to be rapidly passed from the ER Hsp70 chaperone BiP to the lectin chaperones, further underscoring their central role in controlling protein homeostasis in the secretory pathway (Helenius and Hammond, 1994).

N-glycan trimming to an unglucosylated glycoform by glucosidase II supports substrate release from the lectin chaperones. At this stage, if the protein folds properly, it is packaged into COPII vesicles for anterograde trafficking (Barlowe and Helenius, 2016). Alternatively, substrates that are evaluated to be non-native are directed for rebinding to the lectin chaperones by the protein folding sensor UDP-glucose: glycoprotein glucosyltransferase 1 (UGGT1) that reglucosylates immature or misfolded proteins (Helenius, 1994; Sousa and Parodi, 1995). Since UGGT1 directs the actions of this versatile lectin chaperone system and thereby controls protein trafficking through the ER, it acts as a key gatekeeper of the early secretory pathway. Therefore, it is vital to understand the activity of UGGT1 and the scope of substrates it modifies.

Our current knowledge of the activity of UGGT1 relies largely on studies using purified components. UGGT1 was found to recognize non-native or near-native glycoproteins with exposed hydrophobic regions using *in vitro* approaches where the modification of glycopeptides, engineered or model substrates by purified UGGT1 was monitored (Ritter and Helenius, 2000; Taylor et al., 2003; Caramelo et al., 2004). Recent crystal structures of fungal UGGT1 have shown that it possesses a central, hydrophobic cavity in its protein sensing domain, which may support hydrophobic-based interactions for substrate selection (Roversi et al., 2017; Satoh et al., 2017).

Cell-based studies of UGGT1 have relied on the overexpression of cellular and viral proteins (Soldà et al., 2007; Pearse et al., 2008; Ferris et al., 2013; Tannous et al., 2015). *Uggt1* knockout studies have found that the roles of UGGT1 appear to be substrate specific as UGGT1 can promote, decrease or not affect the interaction between substrates and calnexin (Soldà et al., 2007). Prosaposin, the only known cellular substrate of UGGT1 when expressed at endogenous levels, grossly misfolds in the absence of *Uggt1* and accumulates in aggresome-like structures (Pearse et al., 2010). Work in animals has further emphasized the importance of UGGT1 as the deletion of *Uggt1* in mice is embryonically lethal (Molinari et al., 2005).

UGGT1 has a paralogue, UGGT2, but it has no demonstrated cellular activity (Arnold et al., 2000). Domain swapping experiments have demonstrated that UGGT2 possesses a catalytically active glucosyltransferase domain when appended to the folding sensor domain of UGGT1 (Arnold and Kaufman, 2003). *In vitro* experiments using purified, chemically glycosylated interleukin-8 (IL-8), which is not glycosylated in cells, have found that UGGT2 can glucosylate IL-8 (Takeda et al., 2014). This suggests that UGGT2 may be an additional reglucosylation enzyme or protein folding sensor of the ER.

Unlike the classical ATP-dependent chaperones that directly query the conformation of their substrates (Balchin et al., 2016), binding to the lectin chaperones is dictated by enzymes that covalently modify the substrate (Helenius and Aebi, 2004; Hebert et al., 2014). Rebinding to the carbohydrate-dependent chaperones is initiated by the UGGTs that interrogate the integrity of the structure of the protein. Therefore, the proteome-wide detection of cellular UGGT substrates provides the unprecedented opportunity to identify clients that require multiple rounds of chaperone binding and are more reliant on lectin chaperone binding for proper maturation and sorting. Therefore, we designed a cell-based quantitative glycoproteomics approach to identify high-confidence endogenous substrates of UGGT1 and UGGT2 by the affinity purification of monoglucosylated substrates in CRISPR/Cas9-edited cells. UGGT1 and UGGT2 substrates were found to display multiple features of complex proteins including extended lengths plus large numbers of Cys residues and N-glycans. Specific substrates of either UGGT1 or UGGT2 were also discovered, therefore determining that UGGT2 possessed glucosyltransferase activity and identifying its first natural substrates. UGGT1 demonstrated a slight preference for transmembrane proteins, especially those targeted to the plasma membrane, while UGGT2 modification favored soluble lysosomal proteins. The identification of reglucosylated substrates improves our understanding of their folding and maturation pathways and has implications regarding how folding trajectories may be altered in disease states.

## RESULTS

### Experimental design

To identify the substrates that are most dependent upon persistent calnexin/calreticulin cycle binding, we isolated and identified endogenous substrates for the ER protein folding sensors UGGT1 and UGGT2. As the product of a reglucosylation by the UGGTs is a monoglucosylated N-glycan, the presence of the monoglucosylated glycoform was used as a readout for substrate reglucosylation. N-glycans are originally transferred to nascent glycoproteins containing three glucoses, therefore a monoglucosylated glycan can be generated either through trimming of two glucoses from the nascent N-linked glycan or through reglucosylation by the UGGTs. In order to isolate the reglucosylation step from the trimming process, a gene edited cell line was created that transfers abbreviated unglucosylated N-linked glycans to nascent chains. The N-linked glycosylation pathway in mammalian cells is initiated through the sequential addition of monosaccharides, mediated by the *ALG* (Asn-linked glycosylation) gene products, to a cytosolically exposed dolichol-P-phosphate embedded in the ER membrane (Aebi, 2013; Cherepanova et al., 2016) (Figure 1A). The immature dolichol-P-phosphate precursor is then flipped into the ER lumen and sequential carbohydrate addition is continued by additional ALG proteins. The completed N-glycan (Glc_3_Man_9_GlcNAc_2_) is then appended to an acceptor Asn residue in the sequon Asn-Xxx-Ser/Thr/Cys (where Xxx is not a Pro) by the oligosaccharyl transferase (OST) complex (Cherepanova et al., 2016). Initially, a Chinese Hamster Ovary (CHO) cell line with a defect in *Alg6* was employed to establish the utility of this approach to follow (re)glucosylation (Quellhorst et al., 1999; Cacan et al., 2001; Pearse et al., 2008, 2010; Tannous et al., 2015). As the CHO proteome is poorly curated compared to the human proteome, CRISPR/Cas9 was used to knock-out the *ALG6* gene in HEK293EBNA1-6E cells to provide a cellular system that transferred non-glucosylated glycans (Man_9_GlcNAc_2_) to substrates. In these *ALG6*^−/−^ cells, a monoglucosylated glycan is solely created by the glucosylation by the UGGTs providing a suitable system to follow the glucosylation process (Figure 1B).

**Figure 1.**
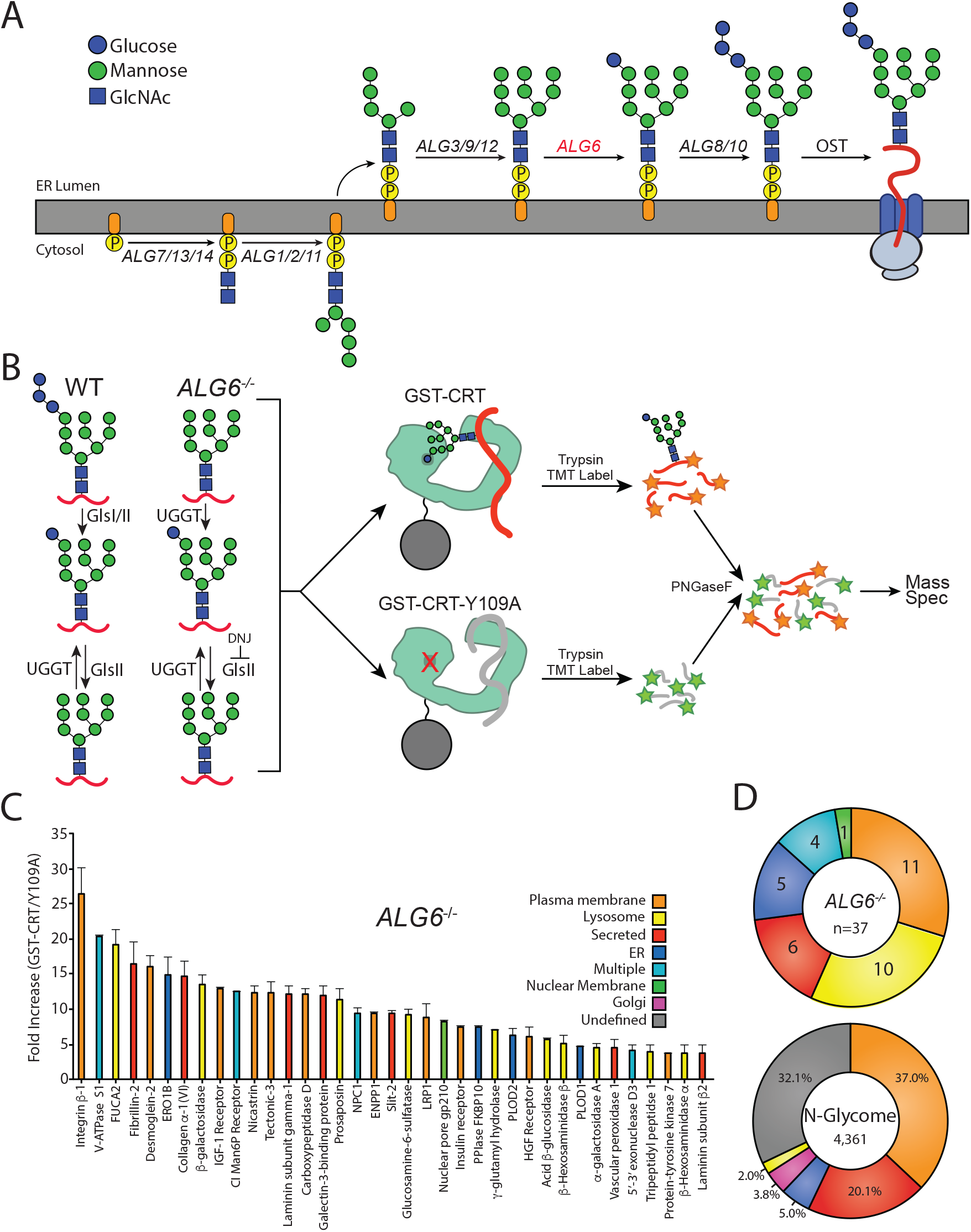
The identification of UGGT1/2 substrates. (A) The pathway of N-glycosylation in eukaryotic cells is depicted. N-glycan synthesis is initiated in the outer ER membrane leaflet on a dolichol-P-phosphate facing the cytoplasm. Flipping of the precursor N-glycan to the ER luminal leaflet and further synthesis steps mediated by ALG proteins leads to eventual transfer of a Glc_3_Man_9_GlcNAc_2_ N-glycan to a substrate by the OST complex. ALG6 (red lettering) catalyzes the transfer of the initial glucose onto the Man_9_ precursor N-glycan. (B) In wild type (WT) cells, a Glc_3_Man_9_GlcNAc_2_ N-glycan is transferred to substrates. Monoglucosylated substrates may therefore occur via trimming by glucosidases I/II (GlsI/II) or reglucosylation by UGGT1/2. In *ALG6*^−/−^ cells, a Man_9_GlcNAc_2_ N-glycan is transferred to substrates. Therefore, monoglucosylated substrates may only occur through reglucosylation by UGGT1/2. DNJ (500 μM) was added to block the trimming of monoglucosylated substrates by GlsII. *ALG6*^−/−^ cells were then lysed and split equally between affinity purifications with either GST-CRT or GST-CRT-Y109A bound to glutathione beads. Affinity-purified samples were then reduced, alkylated, trypsinzed, and labeled with TMT labels. Samples were then deglycosylated with PNGaseF, pooled, and analyzed by mass spectrometry. (C) Substrates were identified by dividing the quantification of the TMT label in the GST-CRT condition for each protein by that of the associated GST-CRT-Y109A condition, yielding the fold increase. Localization as predicted by Uniprot annotation is depicted. A cutoff of three-fold increase was applied. Data is representative of two independent experiments. Error bars represent standard error of the mean (SEM). (D) The N-glycome was computationally determined by collecting all proteins annotated to contain an N-glycome by Uniprot. Annotated localization information was then used to computationally determine the localization distribution of the N-glycome as well as the identified UGGT substrates.

To aid in substrate identification, an inhibitor of glucosidases I and II, deoxynojirimycin (DNJ), was added 1 hr prior to cell lysis to block glucose trimming and trap monoglucosylated products. Monoglucosylated substrates were then isolated by affinity purification using recombinant glutathione S-transferase-calreticulin (GST-CRT), as calreticulin binds monoglucosylated proteins. To account for non-specific binding, a lectin-deficient construct (GST-CRT-Y109A) was used as an affinity purification control (Kapoor et al., 2004). Affinity purified substrates were reduced, alkylated, and trypsin digested. The resulting peptides were labeled with tandem mass tags (TMT) (Rauniyar and Yates, 2014), deglycosylated using PNGaseF, and analyzed by mass spectrometry to identify substrates of the UGGTs. The use of TMT, as well as the control GST-CRT-Y109A affinity purification, allows for robust, quantitative identification of substrates of the UGGTs. The resulting data was analyzed by calculating the fold change in abundance of the TMT associated with proteins identified through affinity purification using wild type GST-CRT over affinity purification using GST-CRT-Y109A. To be considered a UGGT substrate, a cutoff of three-fold (wild type GST-CRT/GST-CRT-Y109A) was applied. This conservative cutoff was set to give a high level of confidence in the identified substrates, as below this cutoff, increasing fractions of non-secretory pathway proteins were found.

### Substrate identification of the UGGTs

In order to determine the cellular substrates of the UGGTs, the above glycoproteomics protocol was followed using *ALG6*^−/−^ cells. A restricted pool of thirty-seven N-linked glycosylated proteins was identified as substrates of the UGGTs (Figure 1C, Supplemental Table 1). Prosaposin, the only previously known endogenous substrate of the UGGTs, was included in this group, supporting the utility of the approach (Pearse et al., 2010). Integrin β-1 showed the most significant fold change (wild type GST-CRT/GST-CRT-Y109A) of ^~^26-fold, indicating there is a large dynamic range of reglucosylation levels.

The cell localizations of UGGT substrates were then determined by using their Uniprot classification. Approximately two thirds of the UGGT substrates are destined for the plasma membrane or lysosomes (Figure 1C and D). Additional substrates are secreted or are resident to the ER or nuclear membrane. Nuclear pore membrane glycoprotein 210 (NUP210) was the only nuclear membrane protein found to be reglucosylated and it is the sole subunit of the nuclear pore that is N-glycosylated (Beck and Hurt, 2016). The nucleus and ER share a contiguous membrane. Proteins targeted to the nuclear membrane are first inserted into the ER membrane, then move laterally to the nuclear membrane (Katta et al., 2014). Four proteins were designated as ‘multiple localizations’ including cation-independent mannose-6-phosphate receptor (CI-M6PR), which traffics between the Golgi, lysosome and plasma membrane (Dell’Angelica and Payne, 2001).

To distinguish the general pool of substrates that the UGGTs are expected to be exposed to, N-glycosylated proteins of the secretory pathway proteome (N-glycome) were computationally defined (Supplemental Table 2). The N-glycome is comprised of proteins that are targeted to the ER either for residency in the secretory/endocytic pathways or for trafficking to the plasma membrane or for secretion. The reviewed UniprotKB *H. sapiens* proteome (20,353 total proteins) was queried to identify all proteins annotated as N-glycosylated, resulting in a set of 4,520 proteins. This set was then curated to remove proteins predicted to be mitochondrial, contain less than 50 amino acids or redundant isoforms. The resulting N-glycome contained 4,361 proteins, predicting ^~^21% of the proteome is N-glycosylated. Comparing UGGT substrates to the N-glycome allows for the characterization of feature preferences of substrates for the UGGTs.

The majority of the N-glycome was either localized to the plasma membrane (37%) or was secreted (20%) according to their Uniprot designations. Smaller fractions of the N-glycome reside in the ER (5%), Golgi (4%) or lysosomes (2%). UGGT substrates are therefore significantly enriched for lysosomal proteins compared to the N-glycome, while all other localizations display a similar distribution to their availability. In total, these results demonstrate the ability to identify substrates of the UGGTs proteomically and suggest that the UGGTs display substrate preferences.

### Determination of UGGT1 and UGGT2 specific substrates

There are two ER glucosyltransferase paralogues, UGGT1 and UGGT2, though currently there is no evidence that UGGT2 acts as a protein sensor or a glucosyltransferase in the cell. Therefore, we sought to determine if UGGT2 has glucosyltransferase activity in the cell, and if so, do these two paralogues have different substrate specificities. To address this concern, GST-CRT affinity purification and TMT mass spectrometry were used to identify substrates of UGGT1 in *ALG6*/*UGGT2*^−/−^ cells and potential UGGT2 substrates in *ALG6*/*UGGT1*^−/−^ cells.

With the *ALG6*/*UGGT2*^−/−^ cells, 66 N-glycosylated proteins were identified as reglucosylation substrates using the three-fold cutoff (GST-CRT/CST-CRT-Y109A) (Figure 2A). Nearly double the number of UGGT1 substrates were identified through this approach compared to using *ALG6*^−/−^ cells where both UGGT1 and UGGT2 were present. This expansion in substrate number is likely due to the ^~^50% increase in expression of UGGT1 in *ALG6*/*UGGT2*^−/−^ cells (Supplemental Figure 1). The substrate demonstrating the most significant fold change (23.5-fold) was CD164, creating a similar dynamic range for reglucosylation to that observed in *ALG6*^−/−^ cells.

**Figure 2.**
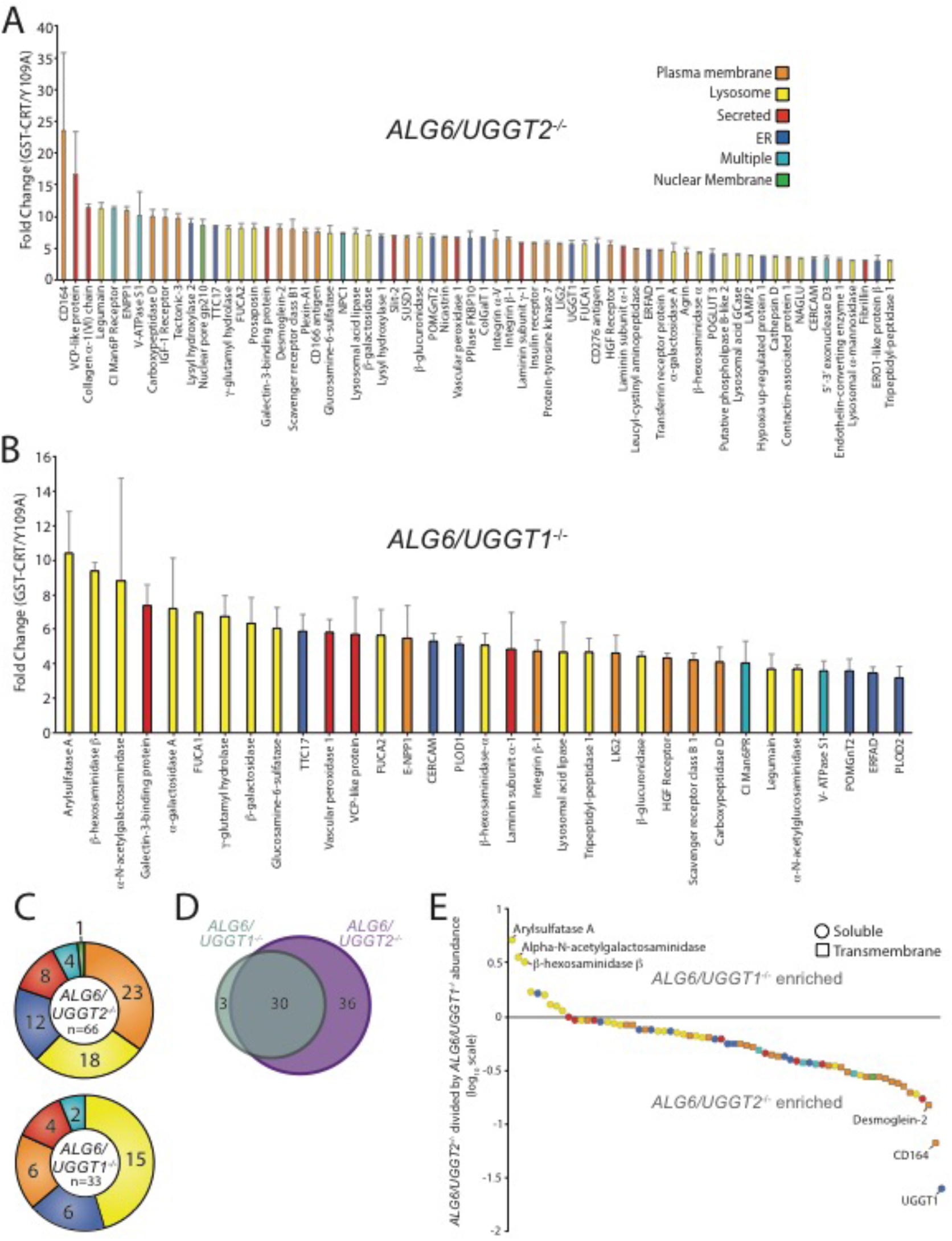
Identification of UGGT1 and UGGT2 specific substrates. (A) Reglucosylation substrates in *ALG6/UGGT2*^−/−^ cells were identified and quantified as previously described in Figure 1. Localizations as annotated by Uniprot are depicted. Data are representative of two independent experiments. Error bars represent SEM. (B) Reglucosylation substrates in *ALG6/UGGT1*^−/−^ cells were identified and quantified as previously above. (C) The distribution of localizations as annotated by Uniprot for reglucosylation substrates identified in both *ALG6/UGGT2*^−/−^ and *ALG6/UGGT1*^−/−^ cells is depicted. (D) The overlap of reglucosylation substrates identified in both *ALG6/UGGT2*^−/−^ cells (purple) and *ALG6/UGGT1*^−/−^ cells (grey) is visualized by a Venn diagram. (E) Reglucosylation substrate enrichment in either *ALG6/UGGT1*^−/−^ or *ALG6/UGGT2*^−/−^ cells is depicted by dividing the TMT quantification for each protein in *ALG6/UGGT2*^−/−^ cells by the associated value in *ALG6/UGGT1*^−/−^ cells on a log_10_ scale. Positive and negative values represent enrichment in *ALG6/UGGT1*^−/−^ and *ALG6/UGGT2*^−/−^ cells, respectively. Localization based on Uniprot annotation is depicted. Proteins are depicted as either circles (soluble) or squares (transmembrane), as annotated by Uniprot.

To identify possible UGGT2 specific substrates, *ALG6/UGGT1*^−/−^ cells were used to isolate UGGT2 modified substrates. Thirty-four proteins passed the three-fold GST-CRT/GST-CRT-Y109A cutoff, with 33 of these proteins predicted to be N-glycosylated and localized to the secretory pathway (Figure 2B). Importantly, this demonstrated for the first time that UGGT2 was a functional glycosyltransferase capable of reglucosylating a range of cellular substrates. The glycoprotein with the most significant fold change was arylsulfatase A (10.4-fold). Notably, 8 of the 9 strongest UGGT2 substrates or, 15 of 33 substrates overall, are lysosomal proteins (Figure 2B and C). While UGGT1 was also observed to engage a significant percentage of lysosomal proteins (27%), 45% of UGGT2 substrates are lysosomal. Both of these percentages are significantly enriched when compared to the N-glycome for which only 2% is comprised of resident lysosome proteins (Figure 1D).

UGGT1 substrates were enriched for plasma membrane localized proteins (35%) when compared to UGGT2 substrates (18%), while plasma membrane proteins were found to compose a similar percent of the N-glycome (37%) compared to UGGT1 substrates. Similar percentages of UGGT1 and UGGT2 substrates localize to the ER (18%), are secreted (12%), or are found in multiple localizations (6%) (Figure 2C). Even though 4% of the N-glycome is composed of Golgi proteins (Figure 1D), neither UGGT1 nor UGGT2 appeared to modify Golgi localized proteins.

The number of UGGT1 substrates was double that of UGGT2 suggesting that UGGT1 carried the main quality control load. Only three out of thirty-three UGGT2 substrates were specific to UGGT2. These three UGGT2 specific substrates included arylsulfatase A, α-N-acetylgalactosaminidase and β-hexosaminidase subunit β (HexB), three soluble lysosomal enzymes (Figure 2D and E). Thirty substrates overlapped between UGGT1 and UGGT2, while thirty-six substrates were found to be specific to UGGT1 (Figure 2D and Supplemental Table 3). The preference for the shared substrates was explored by plotting all proteins identified as a substrate of either glucosyltransferase on a log10 scale of the associated TMT value in *ALG6/UGGT2*^−/−^ cells divided by the values in *ALG6/UGGT1*^−/−^ cells (Figure 2E). Proteins enriched as UGGT2 substrates therefore possess positive values while UGGT1 enriched substrates have negative values.

The three substrates found to be specific to UGGT2 clustered away from all other proteins (Figure 2E at the top left). The remaining UGGT2 enriched substrates, except for one ER localized protein, localized to the lysosome. All the UGGT2 favored substrates were soluble proteins. In contrast, UGGT1 favored proteins were greater in number and displayed a diversity of localizations with a preference for plasma membrane proteins. These results indicate that UGGT2 is a functional glucosyltransferase, which preferentially engages soluble lysosomal proteins while UGGT1 modifies a wider variety of proteins with a preference for plasma membrane and transmembrane domain-containing proteins in general.

### Validation of UGGT substrates

Having identified numerous novel substrates of the UGGTs, a select number of these substrates were tested for reglucosylation to validate the identification approach. Substrates were chosen based on a diversity of topologies, lengths, differences in propensities as UGGT1 or UGGT2 substrates and reagent availability. Monoglucosylated substrates were affinity isolated from *ALG6*^−/−^, *ALG6*/*UGGT1^−/−^*, *ALG6*/*UGGT2*^−/−^ and *ALG6*/*UGGT1*/*UGGT2*^−/−^ cells using GST-CRT compared to CST-CRT-Y109A. Substrates were then identified by immunoblotting with the percent reglucosylation determined by subtracting the amount of protein bound by GST-CRT-Y109A from that of GST-CRT, divided by the total amount of substrate present in the whole cell lysate, and multiplying by 100.

CI-M6PR and insulin-like growth factor type 1 receptor (IGF-1R) are both large type I membrane protein that possess multiple N-glycosylation sites (Figure 3D and H). Overall 10% of CI-M6PR was reglucosylated in *ALG6*^−/−^ cells (Figure 3B). The modification level of CI-M6PR was significantly reduced in *ALG6*/*UGGT1*^−/−^, but not *ALG6*/*UGGT2*^−/−^ cells. As a control, reglucosylation was not observed in *ALG6*/*UGGT1*/*UGGT2*^−/−^ cells. A similar profile was observed for IGF-1R where reglucosylation levels reached 12% in *ALG6*/*UGGT2*^−/−^ cells (Figure 3E-G). Altogether, these findings were consistent with the quantitative glycoproteomics isobaric labeling results (Figure 3C and G), confirming that CI-M6PR and IGF-1R are efficient substrates of UGGT1.

**Figure 3.**
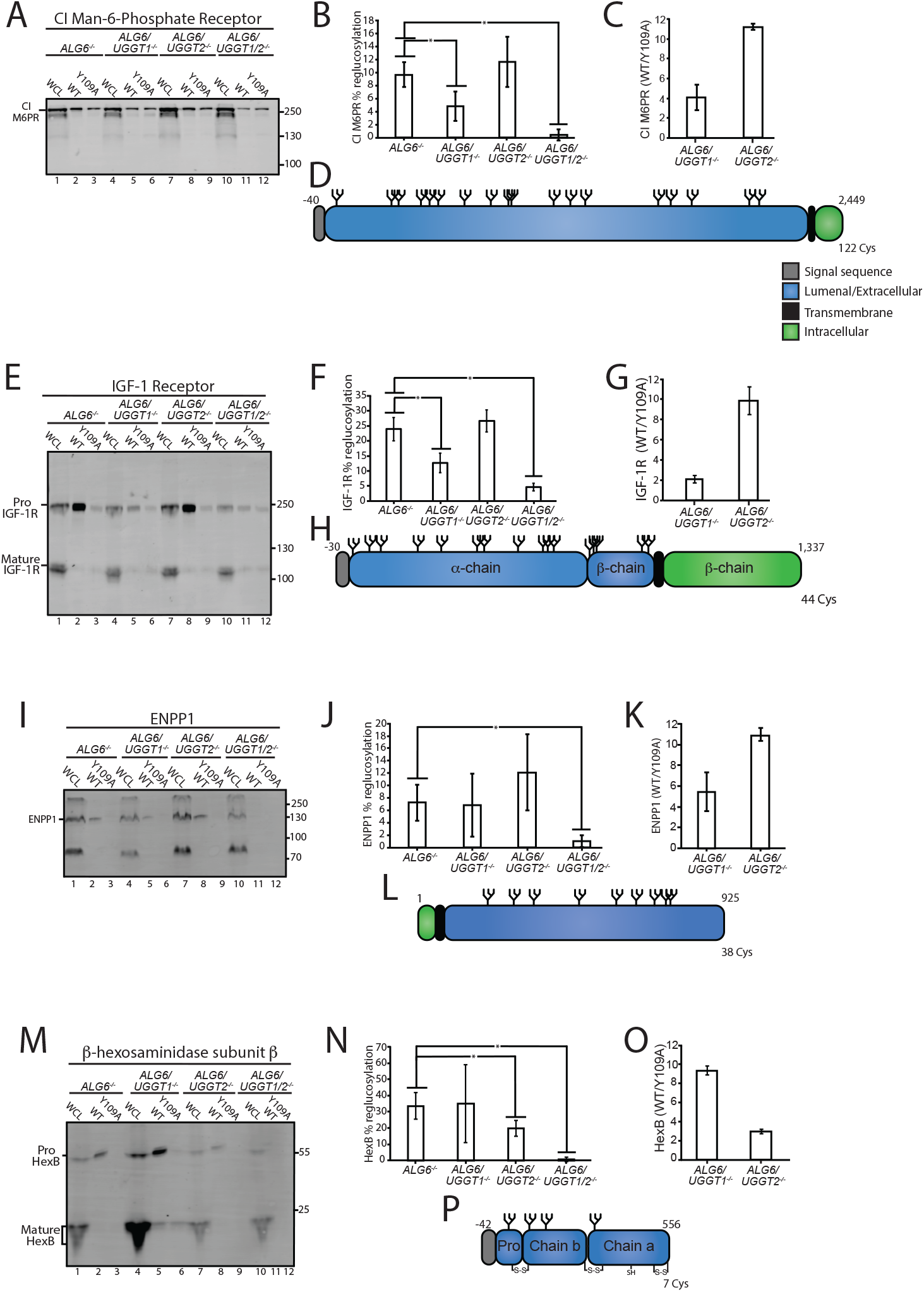
Validation of select reglucosylation substrates. (A) The designated cell lines were lysed and split into whole cell lysate (WCL, 10%) or affinity purification by GST-CRT-WT or GST-CRT-Y109A and imaged by immunoblotting against the CI Man-6-Phosphate receptor. Data is representative of three independent experiments with quantification shown in panel B. Quantifications were calculated by subtracting the value of protein in the Y109A lane from the value of protein in the associated WT lane, divided by the value of protein in the associated WCL lane. Error bars represent the standard deviation. Asterisks denote a p-value of less than 0.05 (C) TMT mass spectrometry quantification of CI Man-6-Phosphate receptor reglucosylation from *ALG6/UGGT1*^−/−^ cells (Figure 2B) and *ALG6/UGGT2*^−/−^ cells (Figure 2A). (D) Cartoon representation of CI Man-6-Phosphate receptor with N-glycans (branched structures), the signal sequence (grey), luminal/extracellular domain (blue), transmembrane domain (black) and intracellular domain (green) depicted. Number of amino acids and Cys residues are indicated. (E) Reglucosylation of IGF-1R, conducted as previously described above. Pro IGF-1R and mature IGF-1R are both observed due to proteolytic processing. Data are representative of three independent experiments with quantification displayed in F. (G) TMT mass spectrometry quantification of IGF-1R from Figure 2A and B, as previously described. (H) Cartoon depiction of IGF-1R. (I) The reglucosylation of ENPP1 shown with quantification displayed in J. (K) TMT mass spectrometry quantification of ENPP1 from Figure 2A and B with cartoon depiction of ENPP1 in L. (M) Reglucosylation of β-hexosaminidase subunit β, conducted as previously described with quantifications displayed in N and TMT mass spectrometry quantification of β-hexosaminidase subunit β from Figure 2A and B in O with a cartoon depicting β-hexosaminidase subunit β in P.

Next, the reglucosylation of the type II membrane protein, ectonucleotide pyrophosphatase/phosphodiesterase family member 1 (ENPP1) was analyzed (Figure 3L). ENPP1 was found to be reglucosylated at similar levels in *ALG6*^−/−^ (7%) and *ALG6*/*UGGT1*^−/−^ (7%) cells. In *ALG6*/*UGGT2*^−/−^ cells, reglucosylation increased to 12%, while in *ALG6*/*UGGT1*/*UGGT2*^−/−^ cells reglucosylation decreased to 1% (Figure 3I and J). These results suggest that ENPP1 can be reglucosylated by both UGGT1 and UGGT2, with a slight preference for UGGT1, supporting the TMT mass spectrometry results (Figure 3K).

The reglucosylation of the smaller soluble lysosomal protein, HexB, was also tested (Figure 3M-P). HexB is processed into three disulfide-bonded chains in the lysosome (Mahuran et al., 1988). Only immature or ER localized proHexB was affinity purified by GST-CRT (Figure 3M, lanes 2, 5, 8 and 11). HexB was reglucosylated at 34% in *ALG6*^−/−^ cells (Figure 3N). No significant change in glucosylation levels were observed when UGGT1 was also knocked out (35%). However, a reduction to 20% reglucosylation of HexB was observed in *ALG6*/*UGGT2*^−/−^ cells, and complete loss of reglucosylation was observed in *ALG6*/*UGGT1*/*UGGT2*^−/−^ cells. *ALG6*/*UGGT1*^−/−^ cells consistently displayed increased levels of expression of HexB (Figure 3M, lane 4), which was supported by RNAseq data (Supplemental Figure 2B). These results confirm the mass spectrometry results, which showed HexB to be a favored substrate of UGGT2 (Fig 3O). It is also notable that HexB, as the first validated substrate of UGGT2, is highly reglucosylated. Taken together, these results demonstrate that the mass spectrometry screen accurately identified substrates of the UGGTs, as well as differentiated between substrates specific to either UGGT1 or UGGT2.

### Analysis of UGGT substrates

To investigate the properties of the substrates modified by the UGGTs and identify potential types of proteins UGGT1 and UGGT2 modify, a systematic analysis of the substrates of the UGGTs was performed and compared to the general properties of the N-glycome. All characteristics were analyzed using UniprotKB annotations. Initially, the length of substrates was compared to the N-glycome. The N-glycome ranged widely in size, from elabela (54 amino acids) to mucin-16 (14,507 amino acids). The overall amino acid distribution of the N-glycome was significantly shifted smaller compared to the size of UGGT substrates (Figure 4A). The median size of the N-glycome was 443 amino acids, compared to 737 for UGGT substrates found in *ALG6*^−/−^ cells. Substrates of both UGGT1 (718 amino acid median) and UGGT2 (585 amino acids) are significantly larger when compared to the N-glycome. This increase in length may lead to more complex folding trajectories, requiring increased engagement with the lectin chaperones for efficient maturation.

**Figure 4.**
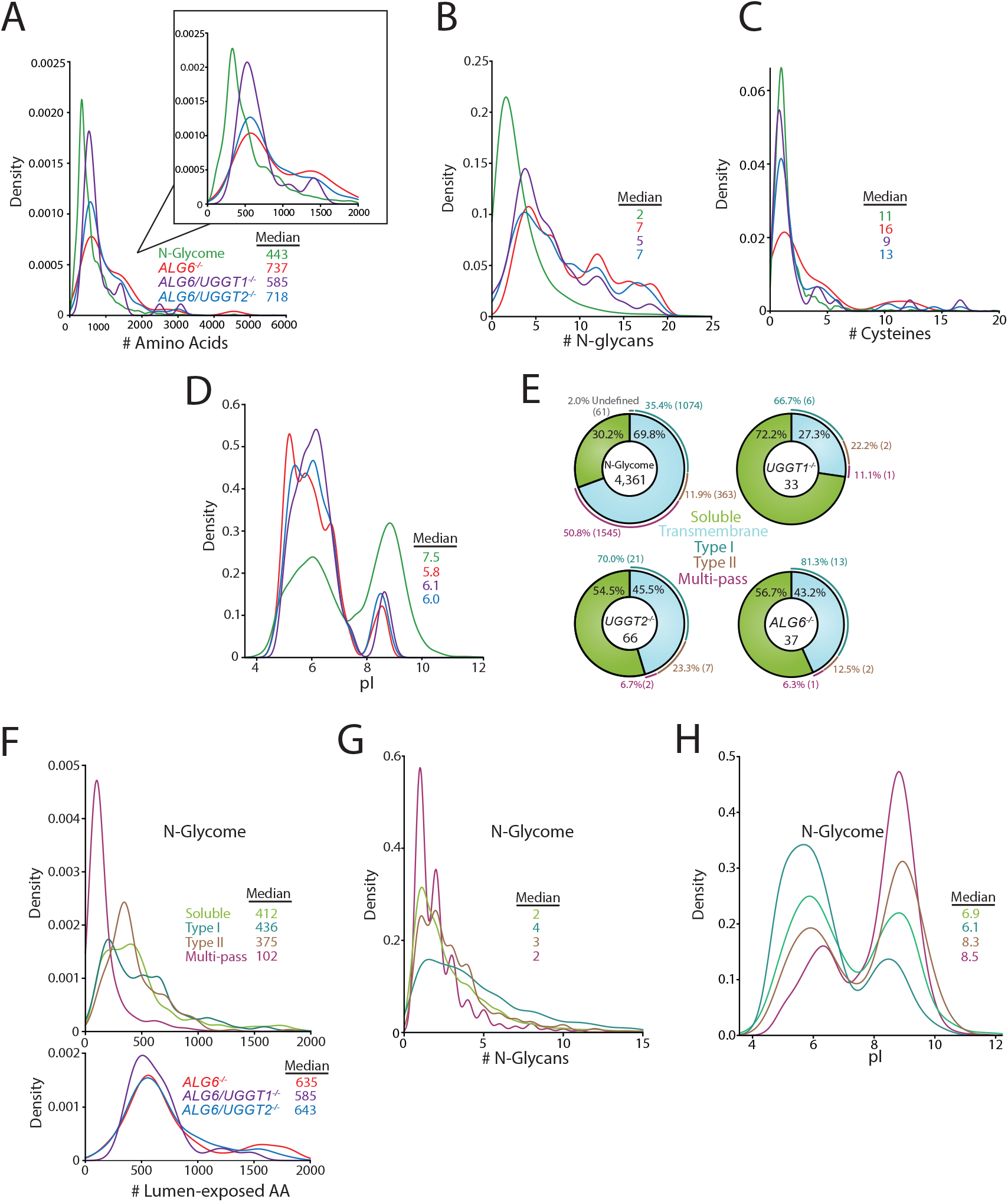
Analysis of substrates of the UGGTs and the N-glycome. (A) Amino acid lengths of each protein in the indicated datasets was visualized by density plot, with the total area under the curve integrated to 1. Amino acid number was obtained via Uniprot annotation. All density plots were generated using R and the ggplot package. (B) The number of N-glycans (B) or Cys residues (C) for each protein in the indicated datasets was visualized by density plot with the numbers determined using their Uniprot annotation. (D) The isoelectric point (pI) values for each protein in the indicated datasets was visualized by density plot. The pI values were obtained via ExPASy theoretical pI prediction. (E) The computationally predicted N-glycome and the indicated reglucosylation substrates were determined as either soluble or transmembrane using Uniprot annotations. The transmembrane portion of each dataset was then analyzed for type I, type II, or multi-pass topology using the associated Uniprot annotation. Proteins which were annotated by Uniprot as transmembrane but lacked topology information were labelled as undefined. (F) The computationally determined N-glycome was separated into soluble, type I, type II, and multi-pass transmembrane proteins using Uniprot annotations. Luminally exposed amino acids were computationally determined using Uniprot annotations for each subset of the N-glycome and each indicated reglucosylation substrate dataset. The resulting data was visualized by density plot. (G) The indicated N-glycome subsets were analyzed for N-glycan content using Uniprot annotation and visualized by density plot, as described. (H) The indicated N-glycome subsets were analyzed for predicted pI using ExPASy theoretical pI prediction and visualized by density plot.

The distribution of the number of N-glycans possessed by the N-glycome (median of 2 glycans per glycoprotein) was also shifted significantly smaller than that of UGGT1 (7 glycans) or UGGT2 (5 glycans) substrates (Figure 4B). All the UGGT substrates displayed both a larger shifted peak and a prominent extended shoulder compared to the N-glycome. Despite the identification of UGGT1 and UGGT2 substrates generally containing high numbers of N-glycans, multiple substrates possessed as few as two N-glycans, suggesting that the experimental approach did not require a high number of monoglucosylated glycans for GST-CRT affinity isolation.

The ER maintains an oxidizing environment that supports the formation of disulfide bonds. Complex folding pathways can involve the engagement of oxidoreductases, such as the calnexin/calreticulin-associated oxidoreductase ERp57, to catalyze disulfide bond formation and isomerization (Margittai and Sitia, 2011; Kozlov and Gehring, 2020). The most common number of Cys residues in proteins identified as UGGT substrates was 2, which was similar to the N-glycome Cys content (Figure 4C). However, there are variations in the median number of Cys residues as for the N-glycome it is 11, which is smaller than that found in *ALG6*^−/−^ cells (16 Cys), and for UGGT1 substrates observed in *ALG6/UGGT2*^−/−^ cells (13 Cys). In contrast, a median of 9 Cys was observed for UGGT2 substrates. Therefore, UGGT1 appears to display a slight preference for proteins with high Cys content, when compared to the N-glycome and UGGT2 substrates.

UGGT1 or UGGT2 substrates displayed similar pI distributions with pIs predominantly near a pH of 6.0, while a second smaller peak centered around a pH of 8.5. Interestingly, a pronounced valley was observed at pH 7.9 under all conditions, presumably due to the instability of proteins with pIs of a similar pH to that of the ER. The N-glycome displayed a more bimodal distribution with significant population of both acidic and basic pIs (Figure 4D). These results suggest that both UGGT1 and UGGT2 preferentially engage proteins with low pIs.

The predicted topologies of the substrates of the UGGTs and the N-glycome were also analyzed. Approximately 70% of the N-glycome is comprised of membrane proteins, with half of these membrane proteins possessing multiple transmembrane domains, followed by single membrane pass proteins with a type I orientation (a third) with the remainder being type II membrane proteins (Figure 4E). A total of 43% of UGGT substrates in *ALG6*^−/−^ cells contained a transmembrane domain with the vast majority of these substrates having their C-terminus localized to the cytosol in a type I orientation, while two substrates possessed the reverse type II orientation and a single multi-pass membrane substrate (NPC1) was identified. When the UGGTs were considered separately, about half of the UGGT1 substrates (*ALG6/UGGT2*^−/−^ cells) possessed at least one transmembrane domain, with 70% of these membrane proteins being in the type I orientation, a quarter in a type II orientation and two being multi-pass proteins (NPC1 and scavenger receptor class B member 1 (SR-BI)). In contrast to UGGT1, the majority of UGGT2 substrates were soluble proteins (72%) with the breakdown of remaining transmembrane proteins being similar to that of UGGT1 with the majority being type I membrane proteins. The preference of UGGTs for type I transmembrane proteins is likely caused by their larger luminal-exposed domains and N-glycan numbers compared to multi-pass membrane proteins (Figure 4F and G). Notably, substrates of the UGGTs had significantly larger luminal domains than the membrane proteins of the N-glycome, though especially for the multi-pass membrane proteins (Figure 4F). Furthermore, while the pIs of type II and polytopic membrane proteins were bimodal, they were overall more basic, which appears to be a property disfavored by UGGT substrates (Figure 4H). Overall, these results show that UGGT1 efficiently modifies both soluble and membrane associated proteins, while UGGT2 strongly favors soluble substrates.

### Efficient IGF-1R trafficking requires lectin chaperone engagement

A number of natural substrates of the UGGTs were identified using a glycoproteomics approach with gene edited cell lines. As reglucosylation by the UGGTs can direct multiple rounds of lectin chaperone binding, the necessity for reglucosylation to support the efficient maturation of a reglucosylated substrate was investigated. IGF-1R is proteolytically processed in the *trans*-Golgi by proprotein convertases including furin, facilitating the monitoring of IGF-1R trafficking from the ER to the Golgi (Lehmann et al., 1998). The requirement for lectin chaperone binding and reglucosylation to aid IGF-1R trafficking was analyzed.

Initially, cells were treated without or with the inhibitor of α-glucosidases I and II, DNJ, to accumulate IGF-1R in the triglucosylated state to bypass entry into the calnexin/calreticulin binding cycle (Helenius and Hammond, 1994; Hebert et al., 1995). At steady state as probed by immunoblotting of cell lysates, IGF-1R accumulated in the ER localized pro form relative to the mature form after DNJ treatment (Figure 5A), resulting in a 19% decrease in the level of the *trans*-Golgi processed mature protein (Figure 5B). This indicated that the lectin chaperone binding cycle helps support efficient IGF-1R trafficking.

**Figure 5.**
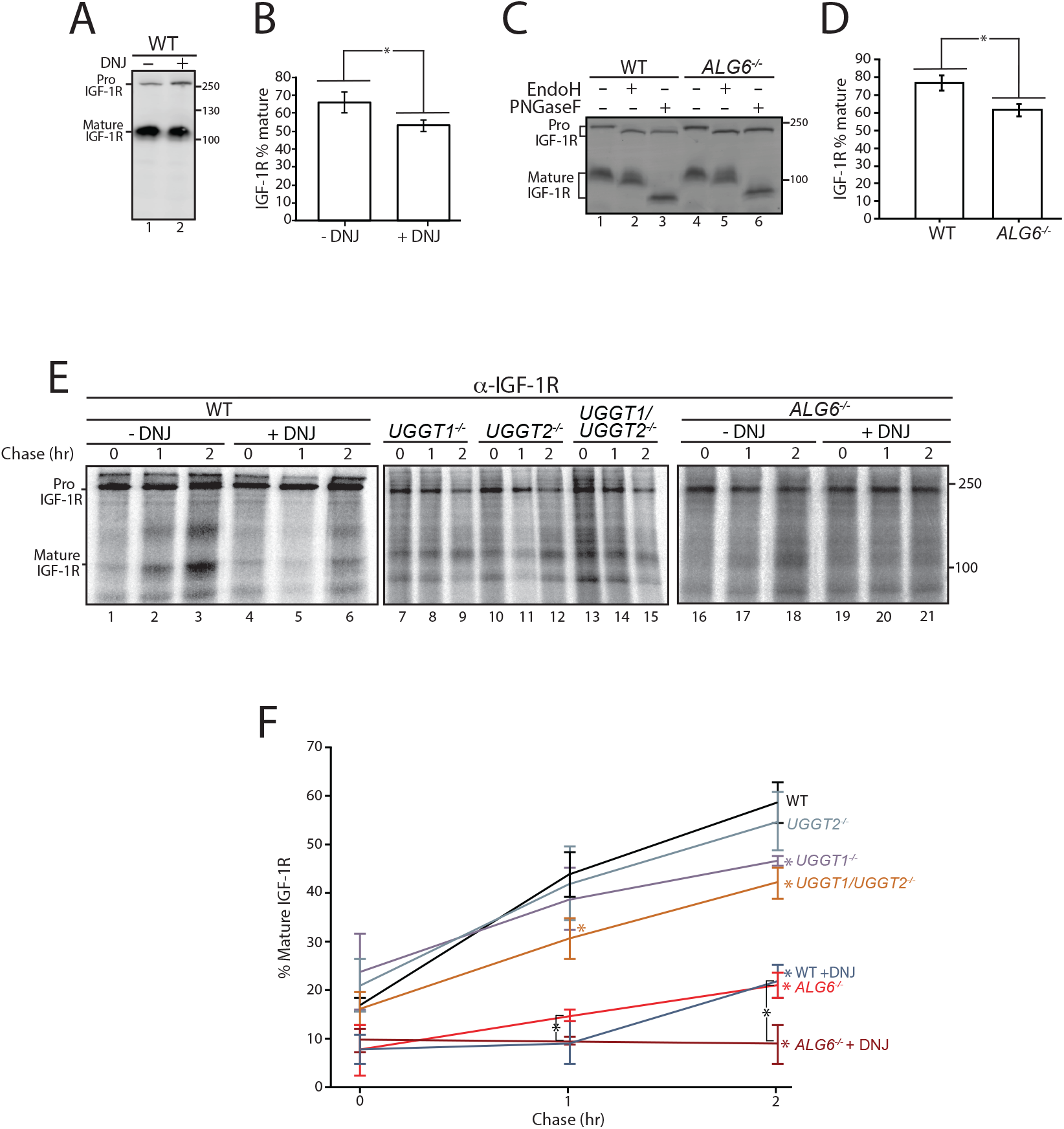
Calnexin/calreticulin cycle role for IGF-1R trafficking. (A) WT HEK293-EBNA1-6E cells treated without or with DNJ (500 μM) for 12-hr were lysed and WCL samples were resolved by reducing 9% SDS-PAGE and imaged by immunoblotting against IGF-1R. Data are representative of three independent experiments with quantification shown in B. Percent of IGF-1R mature was calculated by dividing the amount of mature protein by the total protein in each lane. Errors bars represent standard deviation. Asterisk denotes a p-value of less than 0.05 (C) The indicated cell lines were lysed in RIPA buffer. Samples were split evenly between non-treated and PNGaseF or EndoH treated. Samples were visualized by immunoblotting against IGF-1R and data are representative of three independent experiments with quantification displayed in D. (E) Indicated cells were treated without or with DNJ, pulsed with [^35^S]-Met/Cys for 1-hr and chased for the indicated times. Cells were lysed and samples were immunoprecipitated using anti-β IGF-1R antibody and resolved by reducing SDS-PAGE and imaged by autoradiography. Data are representative of three independent experiments with quantification shown in F

There are two modes for engaging the lectin chaperone cycle: initial binding, which can potentially commence co-translationally for glycoproteins such as IGF-1R that have N-glycans located at their N-terminus through their trimming of the terminal two glucoses by glucosidases I and II; or by rebinding, which is directed by the reglucosylation of unglucosylated species by the UGGTs (Parodi and Caramelo, 2015; Lamriben et al., 2016). The contribution of each mode of monoglucose generation for the proper trafficking of IGF-1R was analyzed.

IGF-1R maturation was investigated in *ALG6*^−/−^ cells as in these cells the N-glycan transferred to the nascent substrate is non-glucosylated, leading to a lack of initial glucosidase trimming mediated lectin chaperone binding. Reglucosylation by the UGGTs is required for lectin chaperone binding in *ALG6*^−/−^ cells. Similar to DNJ treatment in wild type cells, *ALG6*^−/−^ cells demonstrate a 20% decrease in mature IGF-1R relative to the pro form at steady state (Figure 5C, lanes 1 and 3, and Figure 5D). As hypoglycosylation can occur in a substrate dependent manner in *ALG6*^−/−^ cells (Shrimal and Gilmore, 2015), the mobility of IGF-1R with and without N-glycans (PNGase F treated) was monitored by comparing the mobility of IGF-1R by SDS-PAGE and immunoblotting of wild type and *ALG6*^−/−^ cell lysates. IGF-1R, and similarly, CI-M6PR, appeared to be fully glycosylated, while HexB migrated faster when synthesized in *ALG6*^−/−^ cells likely due to hypoglycosylation (Figure 5C and Supplemental Figure 3C and D). To confirm that the pro form of IGF-1R represented ER localized protein rather than protein trafficked out of the ER but not processed by proprotein convertases, IGF-1R from wild-type and *ALG6*^−/−^ cells was treated with the endoglycosidase EndoH. As EndoH cleaves high-mannose glycans which are preferentially present in the ER or early Golgi, an increase in mobility by SDS-PAGE suggests ER localization. In both wild-type and *ALG6*^−/−^ cells, Pro IGF-1R was found to be EndoH sensitive, while mature IGF-1R was found to be largely EndoH resistant (Figure 5C, lanes 2 and 5), suggesting the accumulation of pro IGF-1R in *ALG6*^−/−^ cells represents impaired ER trafficking rather than impaired processing in the *trans*-Golgi. Altogether, these steady state results suggest that lectin chaperone binding is important for efficient IGF-1R maturation.

As steady state results can be impacted by changes in protein synthesis and turnover, a radioactive pulse-chase approach was used to follow protein synthesized during a 1 hr [^35^S]-Met/Cys pulse interval followed by chasing for up to 2-hr under non-radioactive conditions. Pulse-chase experiments are generally performed with overexpressed tag constructs to accumulate and isolate sufficient protein for monitoring. Here, endogenous IGF-1R was isolated by immunoprecipitation with anti-IGF-1R antibodies and analyzed by SDS-PAGE and autoradiography to determine the percent of IGF-1R that was properly processed to its mature form in the *trans*-Golgi. IGF-1R was found to traffic efficiently out of the ER and to the Golgi in wild type cells as 59% of the total protein after a 2-hr chase was mature IGF-1R (Figure 5E, lanes 1-3 and F). When lectin chaperone binding was inhibited by treatment with DNJ, mature IGF-1R was diminished to 22%, underscoring the importance of lectin chaperone binding (Figure 5E, lanes 4-6 and F).

To delineate the contributions of early compared to late lectin chaperone binding, IGF-1R trafficking was followed in gene edited cells that control the methods for lectin chaperone engagement. A single early round of lectin chaperone binding will be permitted in the absence of both UGGTs or rebinding would only be directed by the UGGT present with knockouts of a single UGGT. Alternatively, early lectin chaperone binding as dictated by glucosidase trimming will be absent in the *ALG6*^−/−^ cells where lectin chaperone binding is directed solely through glucosylation by the UGGTs. Monitoring the trafficking of IGF-1R in these cells will allow us to determine the contributions of the different steps in the lectin chaperone binding cycle for proper IGF-1R maturation.

When both UGGTs were absent in *UGGT1/2*^−/−^ cells, the percent of mature IGF-1R after 2 hr of chase decreased to 42%. In agreement with early glycoproteomics and affinity isolation results showing IGF-1R was largely a UGGT1 substrate, UGGT2 knockout alone had little influence on IGF-1R trafficking while the knocking out of UGGT1 supported IGF-1R trafficking similar to the double UGGT deletion (Figure 5E, 7-15 lanes and F). These results support a role for UGGT1 in optimizing IGR-1R trafficking.

To determine the importance of early chaperone binding directed by the glucosidases, IGF-1R trafficking was monitored in *ALG6*^−/−^ cells that support reglucosylation but lack the ability for early binding to the lectin chaperones as directed by glucosidase trimming of the triglucosylated species. In *ALG6*^−/−^ cells, the percent of mature IGF-1R was significantly decreased to 21%, indicative of an important contribution of the initial round of lectin binding, as was suggested by steady state data (Fig 5C). The addition of DNJ to *ALG6*^−/−^ cells would be expected to trap IGF-1R in a monoglucosylated state after glucosylation, allowing the effect of prolonged interaction with the lectin chaperones to be observed. Under this condition, IGF-1R was strongly retained in the ER with no increase observed in the level of mature IGF-1R observed even after 2 hr of chase (Figure 5E, lanes 16-21 and F). Altogether these results demonstrate that while early (glucosidase-mediated) and late (UGGT-mediated) lectin chaperone binding contribute to the efficient trafficking from the ER and subsequent Golgi processing of IGF-1R, early lectin chaperone binding appears to be most critical for supporting proper IGF-1R maturation.

## DISCUSSION

As lectin chaperone binding is directed by the covalent modification of substrates by the UGGTs, the identification of *bona fide* substrates of the UGGTs is central to understand the impact the lectin chaperone network has on cellular homeostasis. Features of proteins alone cannot accurately predict which chaperones will be required for efficient folding and quality control (Adams et al., 2019b). Previous studies involving the UGGTs have focused mainly on the overexpression of biasedly selected substrates or using purified proteins, providing uncertain biological relevance (Ritter and Helenius, 2000; Taylor et al., 2003; Caramelo et al., 2004; Soldà et al., 2007; Pearse et al., 2008; Ferris et al., 2013; Tannous et al., 2015). Here, we used a quantitative glycoproteomics-based strategy to identify seventy-one natural cellular substrates of the UGGTs. When compared to the N-glycome that represents the total population of potential substrates (4,361 N-glycoproteins in human cells), the UGGTs favored the modification of more complex, multidomain proteins with large numbers of N-glycans. These results are in agreement with the common requirement of chaperones for the proper folding of more complex proteins (Balchin et al., 2016, 2020). The lectin chaperone system is part of the robust chaperone network necessary to promote the efficient folding and quality control of substrates and mitigate harmful misfolding events that are associated with a large range of pathologies.

The discovery of 33 UGGT2 cellular substrates provides the first evidence of intact UGGT2 acting as a quality control factor in cells (Figure 2B). Previous work demonstrated that UGGT2 is enzymatically active against chemically engineered glycosylated substrates using purified components or when the catalytic domain of UGGT2 was appended to the folding sensor domain of UGGT1 (Arnold and Kaufman, 2003; Takeda et al., 2014). The lower number of UGGT2 substrates compared to UGGT1 (66 substrates) is likely due, at least in part, to UGGT2 being expressed at a fraction of the level of UGGT1 (^~^4% in HeLa cells (Itzhak et al., 2016)). Of special note is the preference of UGGT2 for lysosomal substrates as 8 of the 9 preferential UGGT2 substrates are lysosomal proteins (Figure 2E). The preferential UGGT2 substrates are all soluble proteins, while half of the preferential UGGT1 substrates contained transmembrane domains indicative of a further preference of UGGT2 for soluble proteins (Figure 2E). Given the preference of UGGT2 for soluble lysosomal proteins, it would be of interest in future studies to examine lysosomes in *UGGT2*^−/−^ cells as a number of the UGGT2 substrates are associated with lysosomal storage diseases including metachromatic leukodystrophy (arylsulfatase A), Sandhoff disease (β-hexosaminidase subunit β) and Schindler disease (α-N-acetylgalactosaminidase) (Mahuran, 1999; Cesani et al., 2016; Ferreira and Gahl, 2017).

UGGT1 serves as the predominant ER glycoprotein quality control sensor. While overall the 66 UGGT1 substrates are evenly distributed between soluble and membrane proteins, the majority of the most efficiently reglucosylated proteins are membrane proteins (Figure 2E). Seventy percent of the membrane proteins modified by UGGT1 are in the type I orientation possessing luminal N-glycosylated domains of significant length. Only two substrates of the UGGTs are multi-pass membrane proteins (NPC1 and SR-BI). In contrast to most polytopic membrane proteins that have little exposure to the ER lumen (Figure 4F), both NPC1 and SR-BI have large heavily glycosylated luminal domains. The enrichment of UGGT1 for transmembrane proteins may be influenced through a weak association with the ER membrane or a general slower and more complex folding process for membrane proteins that provides a longer window for modification.

An important question to ask is what is the basis for the differing substrate specificities of UGGT1 and UGGT2? They display sequence identities that are high within the catalytic domains (83% identical) and lower in their folding sensor domains (49%) (Arnold and Kaufman, 2003). This sequence disparity within the folding sensor domain may drive altered substrate selection. In addition, UGGT1 and UGGT2 may reside in separate subdomains within the ER, which could contribute to substrate accessibility. The CLN6/CLN8 transmembrane complex appears to recognize lysosomal proteins within the ER for COPII packaging in support of a possible mechanism of lysosomal substrate selection (Bajaj et al., 2020). An additional possibility addressed was that the level of expression of the lysosomal proteins identified as UGGT2 substrates may be augmented in *ALG6/UGGT1*^−/−^ cells. However, only the mRNA expression level of β-hexosaminidase subunit β was increased relative to *ALG6*^−/−^ or wild type cells, as supported by immunoblot data (Figure 3M), with the remaining preferential UGGT2 lysosomal substrates displaying no significant change in mRNA expression levels (Supplemental Figure 4). The increased expression of β-hexosaminidase subunit β in *ALG6/UGGT1*^−/−^ cells may be attributed to induction by UPR, as in these cells a slight induction primarily through the ATF6 branch of the UPR was observed (Supplemental Figure 5). Further studies will be required to understand the varying selectivities of the UGGTs.

With some 4,350 possible N-glycosylated proteins as potential UGGT substrates, why were only 71 proteins identified as substrates of the UGGTs? First, many proteins are expected to fold in a chaperone independent manner, especially small, simple proteins. Second, our stringent isolation approach prioritized high quality substrates with at least a 3-fold induction for GST-CRT/GST-CRT-Y109A binding. Third, the profile of reglucosylated substrates is likely cell-type dependent with additional substrates expected to be identified in cell types with heavy secretory pathway loads such as pancreatic cells or hepatocytes, compared to the kidney line used here. Fourth, ^~^1,500 proteins of the N-glycome are multi-pass transmembrane proteins (Figure 4E). This class of protein was strongly de-enriched as substrates of the UGGTs, likely due to their limited luminal exposure and minimal N-glycan content (Figure 4F and G). This reduces the pool of favored substrates by a third. Fifth, the monoglucosylated protein isolation procedure may also be limited by possibly requiring multiple sites of reglucosylation for efficient binding to survive the pulldown protocol. However, multiple substrates with two N-glycans were identified, suggesting heavy glycosylation is not an absolute requirement.

Additionally, protein expression levels are expected to play some role in substrate identification but it does not appear to be a major determining factor as multiple strong substrates were expressed at or below an average protein level for the N-glycome and no correlation between mRNA expression level and the TMT mass spectrometry fold increase for the GST-CRT/GST-CRT-Y109A fraction was observed (Supplemental Figure 2). It would be of interest to determine if proteotoxic stress would increase levels and the range of reglucosylated substrates as both the pool of non-native proteins and the amount of the UPR-induced substrates of the UGGTs would be expected to increase. It is also possible that some proteins identified as substrates of the UGGTs may misfold after missing the first round of calnexin/calreticulin binding in *ALG6*^−/−^ cells and therefore engage the UGGTs more efficiently.

As carbohydrate binding can be dictated initially by glucosidase trimming followed by additional later rounds of binding dictated by UGGT reglucosylation, it is of importance to understand which stage of the binding cycle contributes most significantly to proper protein maturation and cell homeostasis. N-glycans in *Sacchromyces cerevisiae* and other single cell species are transferred post-translationally as they are missing the OST isoform subunit that interacts with the Sec61 translocon and supports early co-translational modification (Ruiz-Canada et al., 2009; Shrimal et al., 2019). A second OST isoform appears in multicellular organisms that is translocon-associated. In addition, reglucosylation activity was first observed in single cell parasites of *Trypanosoma cruzi* where glycans are transferred as Man_9_GlcNAc_2_ moieties thereby bypassing the initial glucosidase initiated binding step observed in metazoans (Parodi and Cazzulo, 1982). These seminal *T. cruzi* studies from Parodi and colleagues that first discovered the (re)glucosylation activity, later attributed to UGGT1, were the inspiration for the development of the experimental *ALG6*^−/−^ system used in this study to isolate substrates of the UGGTs. Conservation analysis of glycosylation and the lectin chaperone pathway suggests that reglucosylation supporting the quality control function of the calnexin cycle evolved prior to its role in assisting in earlier folding events.

Using CRISPR edited cell lines, the contributions of the various steps for chaperone binding engagement for the UGGT1 substrate IGF-1R was experimentally explored as its processing in the Golgi provided a robust Golgi trafficking assay. Furthermore, IGF-1R is a target in cancer biology as it is important for cell growth (Sell et al., 1994; Desbois-Mouthon et al., 2006; Chng et al., 2006; King et al., 2014; Mutgan et al., 2018). When binding to the lectin chaperones was blocked in wild type cells by glucosidase inhibition with DNJ treatment, supporting the production of triglucosylated trapped species, the percent of processed IGF-1R strongly decreased compared to untreated cells, demonstrating a requirement of lectin chaperone engagement for the efficient maturation, trafficking and processing of IGF-1R. In *UGGT1/2*^−/−^ cells, IGF-1R can enter the first round of glucosidase-mediated binding to the lectin chaperones but rebinding directed primarily by UGGT1 mediated reglucosylation cannot occur (Figures 5E, F and 6). This led to a reduced efficiency in the accumulation of mature IGF-1R. The first round of lectin chaperone binding is bypassed in *ALG6*^−/−^ cells as the N-glycans transferred to proteins do not contain glucoses (Figure 6). Therefore, only the rebinding events mediated by reglucosylation take place. More strikingly in *ALG6*^−/−^ cells, this led to a dramatic reduction in IGF-1R processing at a greater level than in *UGGT1/2*^−/−^ cells, indicating the first round of binding to the lectin chaperones was most critical for IGF-1R maturation. The addition of DNJ in *ALG6*^−/−^ cells supported the trapping of reglucosylated side chains and severely reduced Golgi processing, suggesting that reglucosylation-mediated persistent interaction with the lectin chaperones delays IGF-1R exit from the ER.

**Figure 6.**
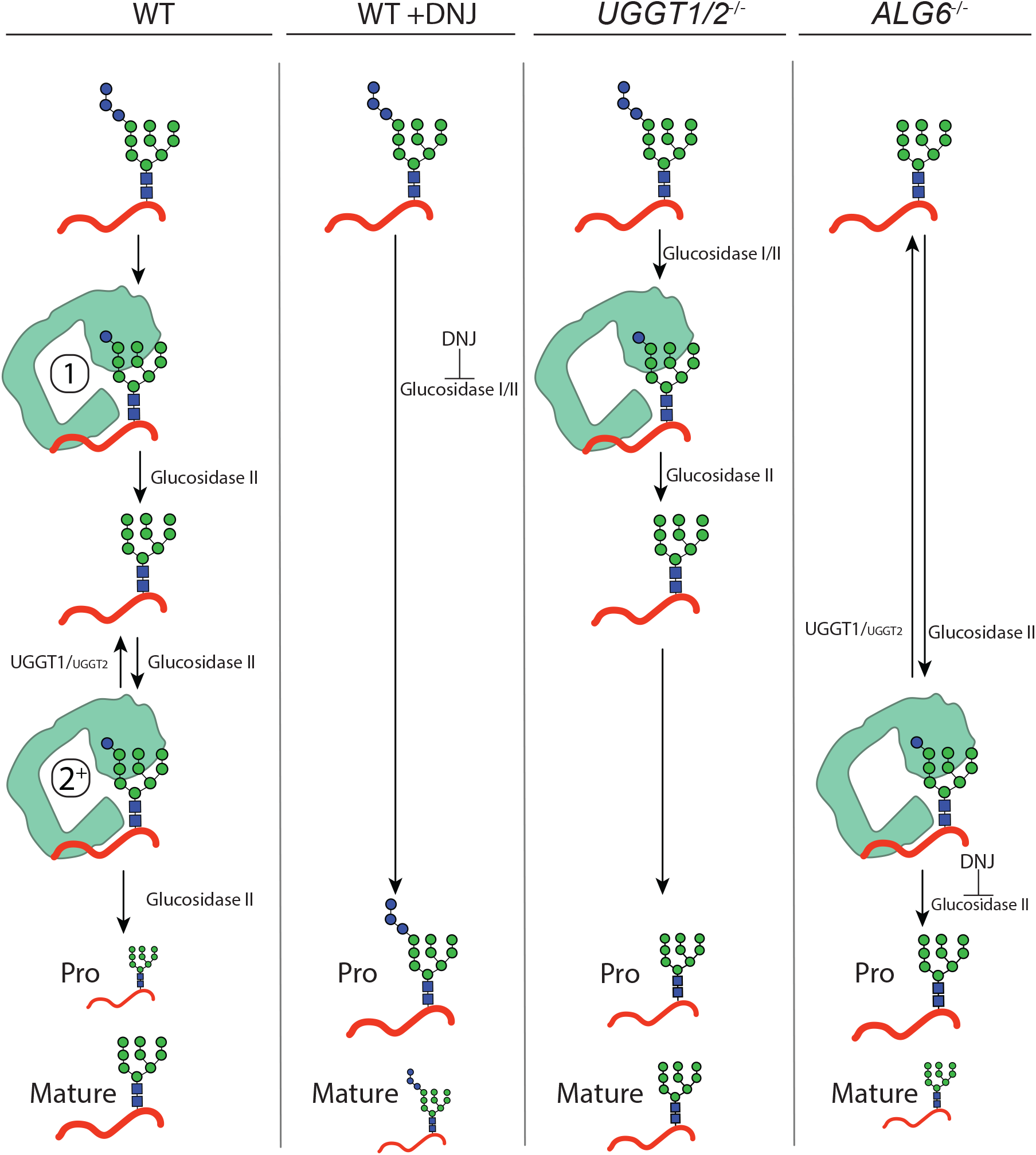
Model for IGF-1R engagement by the lectin chaperone cycle. In WT cells, N-glycans with three terminal glucoses are appended to IGF-1R. Trimming of two terminal glucoses by glucosidases I/II generates a monoglucosylated protein which supports an initial round of interaction with calreticulin (calnexin not shown, denoted by a 1). Trimming of the final glucose by glucosidase II yields a non-glucosylated N-glycan. If recognized as non-native primarily by UGGT1, and to a lesser extent UGGT2, IGF-1R may then be reglucosylated, supporting a second round of interaction with calreticulin (denoted by a 2^+^). Multiple rounds of trimming, reglucosylation and binding to calnexin or calreticulin can occur until proper folding and trafficking. Under this system, IGF-1R is efficiently trafficked from the ER and mature IGF-1R accumulates. When glucosidase I/II activity is inhibited by treatment with DNJ in WT cells, all rounds of binding to the lectin chaperones are ablated and IGF-1R is retained in the ER, yielding primarily pro IGF-1R. In *UGGT1/2*^−/−^ cells, initial binding to calnexin or calreticulin directed by glucosidases I/II trimming is maintained but rebinding via reglucosylation does not occur. Under this system, IGF-1R is inefficiently trafficked from the ER. In *ALG6*^−/−^ cells, N-glycans are transferred without glucoses, eliminating the initial round of binding to calnexin or calreticulin by glucosidases trimming. Only the second round of binding is supported by UGGT1, and to a lesser extend UGGT2, mediated reglucosylation. Upon treatment with DNJ, reglucosylated IGF-1R may persistently interact with the lectin chaperones resulting in ER retention.

Understanding the proteins that interact with or rely on chaperone systems will advance our understanding of protein homeostasis (Houry et al., 1999; Kerner et al., 2005). Large multi-domain proteins such as IGF-1R and many of the other substrates of the UGGTs have apparently evolved to utilize the lectin chaperone system to help direct their complex folding trajectories. The co-evolution of chaperones and their substrates has led to the expansion of the complexity of the proteome for multicellular organisms (Balchin et al., 2016; Rebeaud et al., 2020). The large group of substrates of the UGGTs identified here represents glycoproteins that utilize multiple rounds of lectin chaperone engagement for proper maturation and are likely more prone to misfold under stress. Future studies will determine if this increased vulnerability makes these substrates more susceptible to misfold under disease conditions where cell homeostasis is challenged.

## EXPERIMENTAL METHODS

### Reagents

Antibodies used were: rabbit monoclonal IGF-1 receptor β (D23H3, Cell Signaling), rabbit monoclonal IGF-IIR/CI-M6PR (D3V8C, Cell signaling), rabbit monoclonal BiP (C50B12, Cell Signaling), rabbit monoclonal β-hexosaminidase subunit β (HEXB) (EPR7978, Abcam), rabbit polyclonal ENPP1 (N2C2, Genetex) rabbit polyclonal UGGT1 (GTX66459, Genetex), mouse monoclonal Glyceraldehyde 3-Phosphate (MAB374, Millipore Sigma), IRDye x anti-rabbit secondary (LiCor). All chemicals were purchased from Millipore-Sigma, except where indicated.

### Cell culture

HEK293-EBNA1-6E cells were employed and used as the parental line to create all CRISPR/Cas9 edited lines (Tom et al., 2008). Cells were cultured in DMEM (Sigma) supplemented with certified 10% fetal bovine serum (Gibco) at 37 °C at 5% CO_2_. Cells were tested for the presence of mycoplasma using a universal mycoplasma detection kit (ATCC, Cat # 30-012K).

### CRISPR/Cas9-mediated knock outs

HEK293EBNA1-6E *ALG6*^−/−^, *ALG6/UGGT1*^−/−^, *ALG6/UGGT2*^−/−^, *ALG6/UGGT1/UGGT2*^−/−^, *UGGT1*^−/−^, *UGGT2*^−/−^, and *UGGT1/2*^−/−^ cells were generated via CRISPR/Cas9 using gRNA plasmids gh260, gh172, and gh173, and Cas9-GFP plasmid CAS9PBKS (Lonowski et al., 2017; Narimatsu et al., 2018). Plasmids gh260 (106851), gh172 (106833), gh173 (106834), and CAS9PBKS (68371) were from Addgene. Knock-out cell lines were generated by co-transfecting HEK293-EBNA1-6E cells at 70% confluency in a 10-cm plate with 7 μg of both the associated gRNA and Cas9-GFP plasmid, using a 2.5 μg of PEI per 1 μg of plasmid. Cells were allowed to grow for 48 hr prior to trypsinization and collection. After trypsinization, cells were collected and washed twice with sorting buffer (1% FBS, 1mM EDTA, PBS). Cells were then resuspended in sorting buffer at approximately 1 million cells per ml. Cells were then bulk separated using flow assisted cell sorting based on the top 10% of Cas9-GFP expressing cells (FACS Aria II SORP, Becton Dickinson and Company). Cells were then plated at 5, 10, 20 thousand cells per 10 cm plate in pre-conditioned DMEM media with 20% FBS. Colonies derived from a single cell were isolated using cell cloning cylinders (Bellco Glass), trypsinized from the plate, and further passaged. Knock-outs were confirmed by immunoblotting and staining for UGGT1 or, where antibodies were not available, isolating genomic DNA using a genomic DNA isolation kit (PureLink genomic DNA mini kit, Thermo Fisher), PCR amplification of the genomic DNA region of interest, and insertion of genomic DNA into pcDNA3.1-. Plasmids were then sequenced for conformation (Genewiz).

### GST-CRT purification

The plasmid for pGEX-3X GST-CRT was from Prof. M. Michalak (University of Alberta). pGEX-3X GST-calreticulin-Y109A was generated by site-directed mutagenesis. GST-CRT was expressed in BL21 *E. Coli* cells in LB medium containing ampicillin at 100 μg/ml. Cultures were grown at 37 °C with shaking until an O.D. of A_600_=0.6. Protein expression was then induced by treating cultures with 8.32 mg/L IPTG for 2 hr. Cultures were centrifuged at 3,000 g for 10 min. Cell pellets were lysed with cold lysis buffer (1 mM phenylmethylsulfonyl fluoride, 2% Triton X-100, PBS pH 7.4) and resuspended. Resuspended cells were lysed in a microfluidizer (110L, Microfluidics) at 18,000 psi for two passes. The cell lysate was centrifuged for 40 min at 8,000 g at 4 °C. Lysate was filtered through a 0.45 μm filter. Two ml bed volume glutathione sepharose beads (GE Lifesciences, Cat# GE17-0756-01) per liter of lysate was equilibrated in wash buffer (1% Triton X-100, 1 mM PMSF, PBS pH 7.4), added to cleared lysate, and rotated at 4 °C for 3 hr. Beads were precipitated through centrifugation at 1,000 g for 5 min at 4 °C. The supernatant was aspirated and beads were washed twice in wash buffer with gentle resuspension between washes. One ml of elution buffer (10 mM reduced glutathione, 1 mM PMSF, 50 mM Tris pH 8.5) was added to beads and beads were gently resuspended and allowed to incubate for 5 min at 4 °C. Beads were precipitated by centrifugation at 1,000 g for 5 min 4 °C. The eluate was collected and a total of 6-elutions were collected. Resulting eluate was tested for purity and protein amount on a reducing SDS-PAGE and stained with Imperial protein stain (Thermo Fisher, Cat# 24617). Elutions were then combined and protein concentration was quantified by a Bradford assay (Bio-Rad). Purified protein was then stored at −80 °C in a 20% glycerol PBS buffer at 1 mg/ml.

### GST-CRT isolation and TMT mass spectrometry sample preparation

Five 10 cm plates were seeded with 3.5 million cells and allowed to grow for 48 hr. Cells were treated with N-butyldeoxynojirimycin hydrochloride (DNJ) Cayman Chemicals, Cat # 21065) at 500 μM for 1 hr. Prior to lysis, the media was aspirated and cells were washed once with filter sterilized PBS. Cells were lysed in 1 ml of lysis buffer (20 mM MES, 100 mM NaCl, 30 mM Tris pH 7.5, 0.5% Triton X-100) per plate. Samples were shaken at 4 °C for 5 min and centrifuged at 20,800 g at 4 °C for 5 min. Lysate was pre-cleared with 25 μl bed volume of buffer-equilibrated glutathione beads per 1 ml of lysate under rotation for 1 hr at 25 μl bed volume. Beads were precipitated by centrifugation at 950 g at 4 °C for 5 min. Glutathione beads were pre-incubated with either GST-CRT or GST-CRT-Y109A by equilibrating 25 μl bed volume/pull-down glutathione beads with lysis buffer. Beads were incubated with 100 μg of purified GST-CRT/pull-down under gentle rotation at 4 °C for 3 hr. Beads were centrifuged at 950 g at 4 °C for 5 min and washed twice with lysis buffer. Supernatant was collected and split in half, with one half incubated for 14 hr at 4 °C under gentle rotation with glutathione beads pre-incubated with GST-CRT and the other half under the same conditions with GST-CRT-Y109A.

After incubation with GST-CRT beads, samples were washed once in lysis buffer without protease inhibitors and twice in 100 mM triethylammonium bicarbonate (Thermo Fisher Cat# 90114). After the final wash, samples were incubated with 10 μl of 50 mM DTT (Pierce, Cat# A39255) for 1 hr at room temperature under gentle agitation. Samples were treated with 2 μl of 125 mM iodoacetamide (Pierce, Cat# A39271) and incubated for 20 min under gentle agitation, protected from light. Samples were digested with 5 μg of trypsin (Promega, Cat# V5280) at 37 °C overnight under agitation. Peptide concentration was quantified using a BCA protein quantification kit (Pierce, Cat# 23227). 10plex tandem mass tags (TMT) (Themo Fisher 0.8mg) were resuspended in mass spectrometry grade acetonitrile and was added to digested peptide and incubated for 1-hr at room temp, per manufacturer’s instructions. Labeling was quenched by adding hydroxylamine to 0.25% and incubating for 15 min at room temp. Labeled samples were pooled, treated with 1,000 units of glycerol-free PNGaseF (NEB, Cat# P0705S), and incubated for 2-hr at 37 °C. Samples were cleaned using C18 tips (Pierce, Cat# 87784), and eluted in 75% mass spectrometry grade acetonitrile, 0.1% formic acid (TCI Chemicals). Sample peptide concentration was then quantified using a colorimetric assay (Pierce, Cat# 23275).

### Mass spectrometry data acquisition

An aliquot of each sample equivalent to 3 μg was loaded onto a trap column (Acclaim PepMap 100 pre-column, 75 μm × 2 cm, C18, 3 μm, 100 Å, Thermo Scientific) connected to an analytical column (Acclaim PepMap RSLC column C18 2 μm, 100 Å, 50 cm × 75 μm ID, Thermo Scientific) using the autosampler of an Easy nLC 1000 (Thermo Scientific) with solvent A consisting of 0.1% formic acid in water and solvent B, 0.1% formic acid in acetonitrile. The peptide mixture was gradient eluted into an Orbitrap Fusion mass spectrometer (Thermo Scientific) using a 180 min gradient from 5%-40%B (A: 0.1% formic acid in water, B:0.1% formic acid in acetonitrile) followed by a 20 min column wash with 100% solvent B. The full scan MS was acquired over range 400-1400 m/z with a resolution of 120,000 (@ m/z 200), AGC target of 5e5 charges and a maximum ion time of 100 ms and 2 s cycle time. Data dependent MS/MS scans were acquired in the linear ion trap using CID with a normalized collision energy 35%. For quantitation of scans, synchronous precursor selection was used to select 10 most abundant product ions for subsequent MS^3 using AGC target 5e4 and fragmentation using HCD with NCE 55% and resolution in the Orbitrap 60,000. Dynamic exclusion of each precursor ion for 30 s was employed. Data were analyzed using Proteome Discoverer 2.4.1 (Thermo Scientific).

### Computational determination of the human N-glycome and substrates analyses

The human N-glycome was defined by the total predicted N-glycosylated proteins from the reviewed human proteome from the UniprotKB (accessed 8/10/2020). Both manual and automated curation of the data set was preformed to remove mitochondrial proteins as well as proteins smaller than 50 amino acids from the dataset. All annotations were derived directly from the UniprotKB information and annotations available for these proteins were analyzed in R. Determination of the pI values were performed by the pI/MW tool on the Expasy database.

### Reglucosylation validation assay

Five 10 cm plates were seeded with 3.5 million cells each and allowed to grow for 48 hr. Cells were treated with DNJ at 500 μM for 14 hr. Prior to lysis, the media was aspirated and cells were washed once with filter sterilized PBS. Cells were lysed in 1 ml of MNT (20 mM MES, 100 mM NaCl, 30 mM Tris pH 7.5, 0.5% Triton X-100) with protease inhibitors (50 μM Calpain inhibitor I, 1 μM pepstatin, 10 μg/ml aprotinin, 10 μg/ml leupeptin, 400 μM PMSF) and 20 mM N-ethyl maleimide, shaken vigorously for 5 min at 4 °C, and centrifuged for 5 min at 17,000 g at 4 °C. 50 μl bed volume of glutathione beads was added to each pull-down and incubated for 1 hr at 4 °C under gentle rotation. Beads were then precipitated by centrifugation at 1,000 g for 5 min at 4 °C. Supernatant was collected with 10% used for WCL and the remainder split evenly between GST-CRT and GST-CRT-Y109A conjugated glutathione beads, which were generated as previously described, and incubated for 16 hr at 4 °C under gentle rotation. Beads were precipitated at 1,000 g for 5 min at 4 °C. Supernatant was aspirated and beads were washed twice with lysis buffer without protease inhibitors. Beads were treated with reducing sample buffer (30 mM Tris-HCl pH 6.8, 9% SDS, 15% glycerol, 0.05% bromophenol blue). WCLs were trichloroacetic acid (TCA) precipitated by adding TCA to cell lysate to a final concentration of 10%. Cell lysate was then briefly rotated and allowed to incubate on ice for 15 min before centrifugation at 17,000 g for 10 min at 4 °C. Supernatants were aspirated and washed twice with cold acetone and centrifuged at 17,000 g for 10 min at 4°C. Supernatants were aspirated and the remaining precipitant was allowed to dry for 5 min at room temperature and briefly at 65 °C. Precipitated protein was resuspended in sample buffer. Samples were resolved on a 9% reducing SDS-PAGE and imaged by immunoblotting.

### Metabolic labeling and IGF-1R immunoprecipitation

Two million cells were plated in 6 cm plates and allowed to grow for 40 hr. Cells were pulse labeled for 1 hr with 120 μCi of EasyTag Express^35^S Protein Labeling Mix [^35^S]-Cys/Met (PerkinElmer; Waltham, MA). Immediately after the radioactive pulse, cells were washed with PBS and either lysed in MNT with a protease inhibitor cocktail (Halt protease and phosphatase inhibitor single-use cocktail, Thermo Fisher) and 20 mM NEM, or chased for indicated time using regular growth media. Where indicated, cells were treated with 500 μM DNJ for 30 min prior to [^35^S]-Cys/Met labeling and through the chase. Cell lysates were shaken for 5 min at 4 °C, centrifuged at 17,000 g for 5 min at 4 °C, and the supernatants were collected. Samples were pre-cleared with a 20 μl bed volume of protein-A sepharose beads (GE Healthcare) by end-over-end rotation for 1 hr at 4 °C. The supernatants were collected and incubated with a 30 μl bed volume of protein-A-sepharose beads and 1.5 μl of α-IGF-1 receptor β (D23H3) XP (Cell Signaling) per sample. Samples were washed with MNT without protease inhibitors or NEM and eluted in sample buffer. Samples were then resolved on a 9% reducing SDS-PAGE, imaged using a GE Typhoon FLA 9500 phosphorimager (GE Healthcare), and quantified using ImageJ.

### Glycosylation assay

Three million cells for each indicated cell line were plated in a 10-cm plate and allowed to grow for 48 hr. Cells were lysed in 300 μl RIPA buffer (1% SDS, 1% NP-40, 0.5% sodium deoxycholate, 150 mM NaCl, 50 mM Tris-HCl pH 8.0) with protease inhibitor cocktail and 20 mM NEM. Samples were then sonicated for 20-sec at 40% amplitude (Sonics vibra cell VC130PB), shaken vigorously for 5 min, and centrifuged for 5 min at 17,000 g. 20 μl of the resulting lysate was heated at 95 °C for 5 min, and treated with either 10 μl of PNGaseF or EndoH for 1 hr at 37 °C, according to the manufacturer’s instructions (NEB). Samples were diluted 1:1 into sample buffer and imaged by immunoblotting.

### RNAseq library preparation and Sequencing

Three million cells for each indicated cell line were plated in 10 cm plates and allowed to grow for 48 hr. Cells were then lysed in TRIzol buffer and RNA was isolated using RNA Clean Concentrate Kit with in-column DNase-I treatment (Zymo Research Corp), following manufacturer instructions. The quantity of RNA was assayed on Qubit using RNA BR assay (Life Technologies Corp), and quality was assessed on Agilent 2100 Bioanalyzer using RNA 6000 Nano Assay (Agilent Technologies Inc). Total RNA was used to isolate poly(A) mRNA using NEBNext Poly(A) mRNA Magnetic Isolation Module, and libraries were prepared using NEBNext UltraII Directional RNA Library Prep Kit for Illumina (New England Biolabs) following manufacturer instructions. The quantity of library was assayed using Qubit DNA HS assay (Life Technologies Corp), and quality was analyzed on Bioanalyzer (Agilent Technologies Inc). Libraries were sequenced on Illumina NextSeq 500 platform using NextSeq 500/550 High Output v2 kit (150 cycles) with 76 bp paired-end sequencing chemistry.

Sequence quality was assessed using FastQC (Andrews) and MultiQC (Ewels et al., 2016). Reads were aligned to the hg38 human reference genome using STAR (Dobin et al., 2013). Transcript abundance was quantified using RSEM (Li and Dewey, 2011) and normalized to counts per million (CPM) in R using the edgeR software package (Robinson et al., 2010). Analyses to compare gene expression between cell types was conducted in Excel by finding the average CPM in the pool of genes of interest for the associated cell type and determining the standard deviation away from the average for each gene of interest.

## ACKNOWLEDGEMENTS

This work was supported by the National Institutes of Health under award number (GM086874 (to D.N.H.); and a Chemistry-Biology Interface program training grant (T32GM008515 to B.M.A. and N.P.C.). Mass spectral data were obtained at the University of Massachusetts Mass Spectrometry Center (Director Dr. Steve Eyles). Flow cytometry and RNAseq data were conducted at the University of Massachusetts Flow Cytometry Core Facility (Director Dr. Amy Burnside) and the Genomics Resource Laboratory (Director Dr. Ravi Ranjan), respectively. John Swenson conducted the analysis of raw RNAseq data.

