## Supplemental Figures for "Quantitative glycoproteomics reveals cellular substrate selectivity of the ER protein quality control sensors UGGT1 and UGGT2"

Supplemental Figure 1

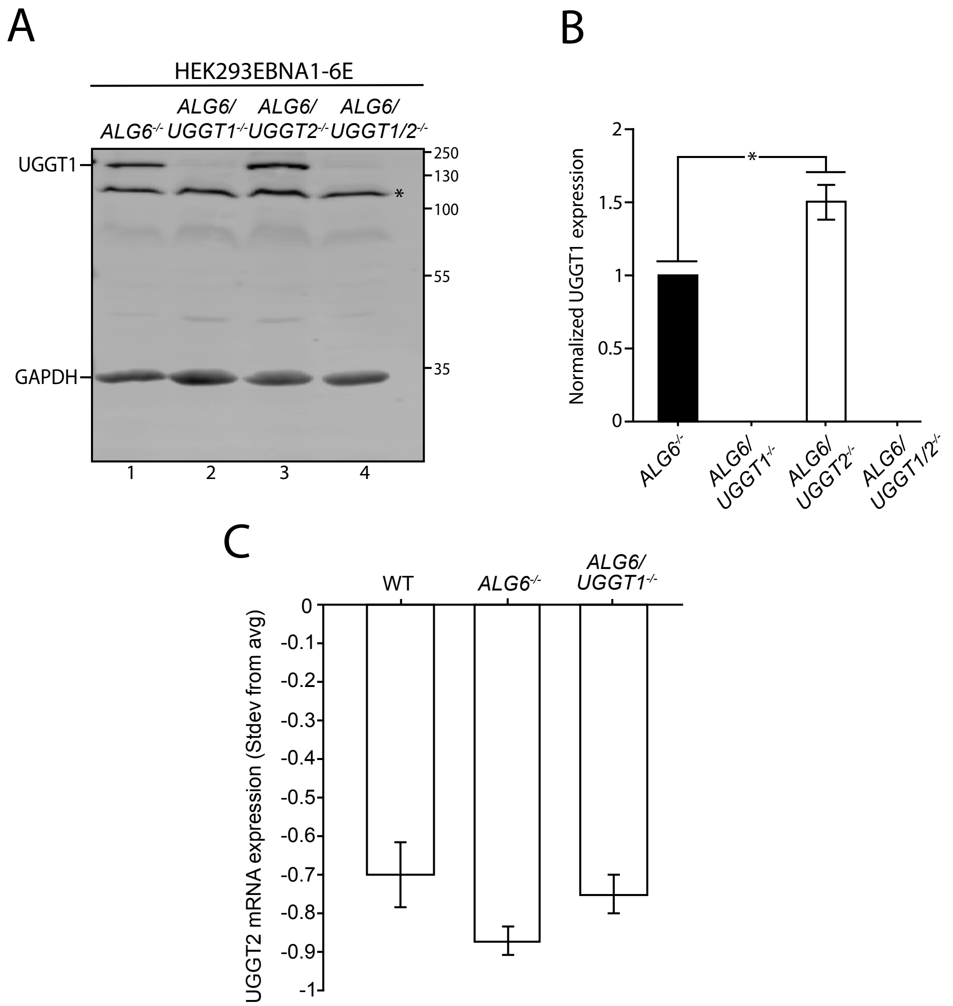


**Supplemental Figure 1. UGGT1 and UGGT2 expression.**

(A) The indicated cells were lysed and whole cell lysates were resolved by SDS-PAGE and imaged by immunoblotting against UGGT1 and GAPDH. Asterisk denotes background band. Data are representative of three independent experiments with quantification shown in B. UGGT1 expression was normalized to that of *ALG6*$^{-/-}$ cells. Error bars represent standard deviation. Asterisk denotes a p-value of less than 0.05. (C) Counts per million of UGGT2 mRNA generated by RNAseq from Supplemental Table 4 was analyzed for the level of *UGGT2* mRNA expression in the indicated cell lines. Counts per million of all genes were averaged and the standard deviation from the average for *UGGT2* mRNA was determined. Error bars represent the standard deviation. Data are representative of three independent experiments.

Supplemental Figure 2

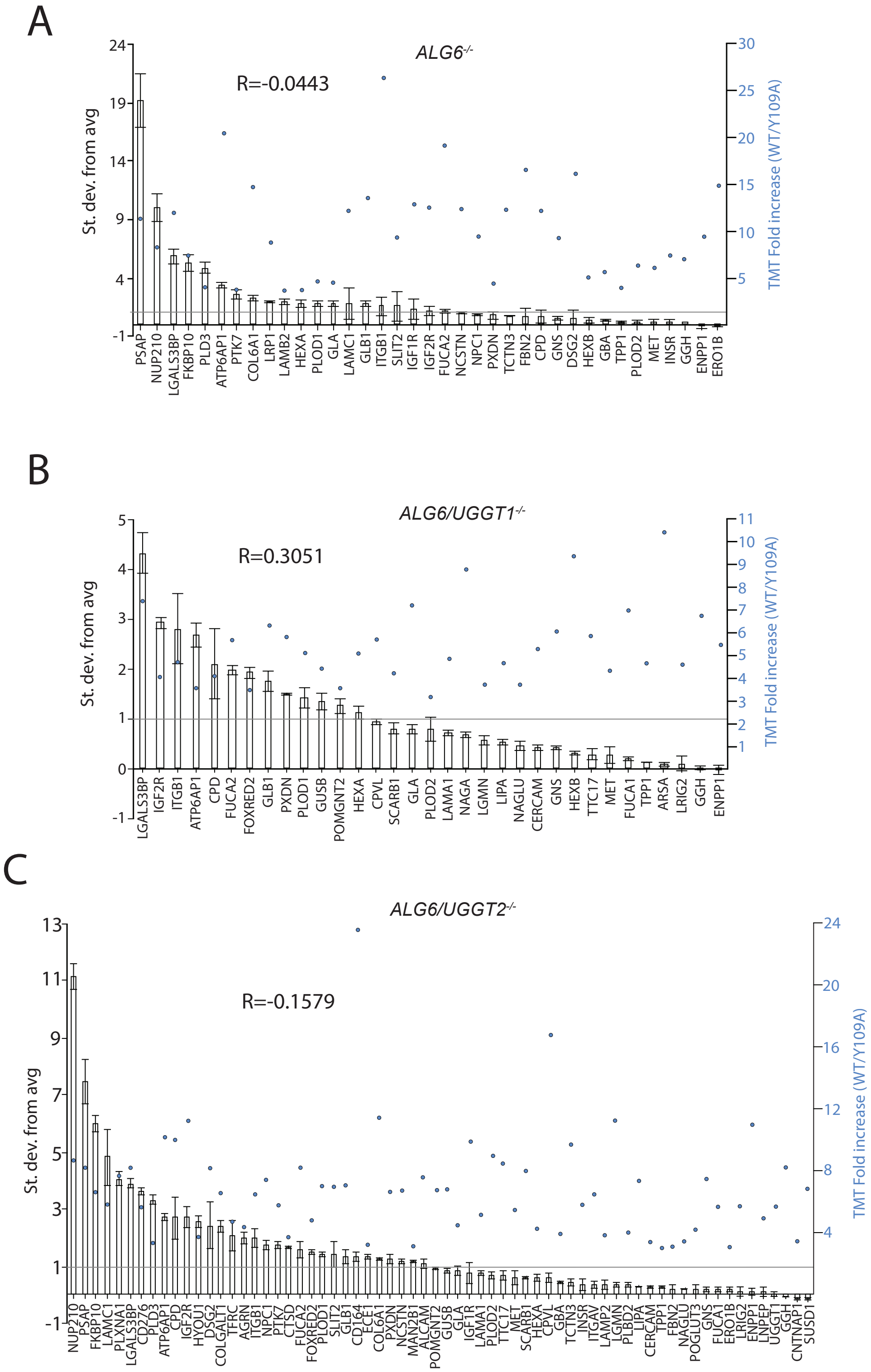

**Supplemental Figure 2. mRNA expression analysis of UGGT1 and UGGT2 substrates.**

(A) Reglucosylated substrates identified in *ALG6*$^{-/-}$ (A), *ALG6/UGGT1*$^{-/-}$ (B) and *ALG6/UGGT2*$^{-/-}$ (C) cells were compared to the average expression for the N-glycome in counts per million. The standard deviation from the average is plotted, with the error bars representing the standard deviation. Blue dots above each gene represent the level of fold increase (GST-CRT/GST-CRT-Y109A) found by TMT mass spectrometry. The Pearson's correlation coefficient (R) between the mRNA expression and TMT mass spectrometry fold increase is shown. Data is representative of three independent experiments.

### Supplemental Figure 3

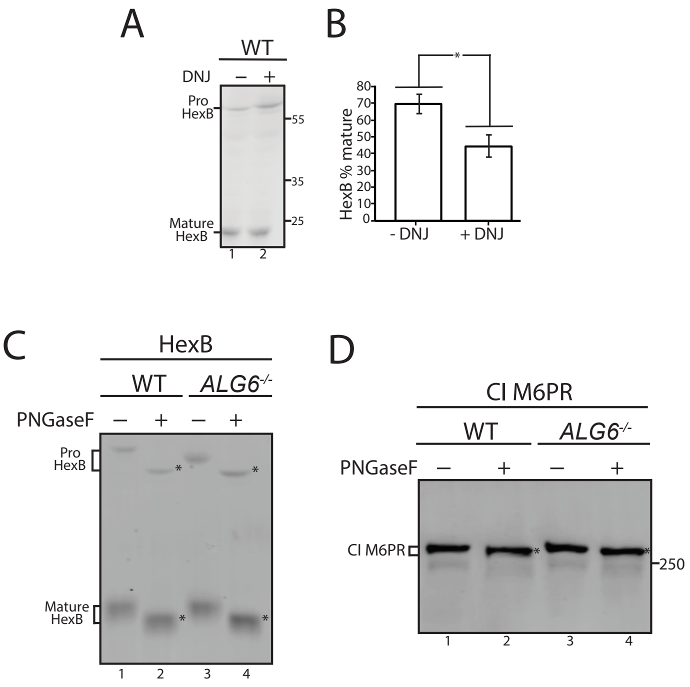


**Supplemental Figure 3. β-hexosaminidase subunit β trafficking and hypoglycosylation**

**and CI M6PR hypoglycosylation.**

(A) Cells treated without or with DNJ for 12 hr, lysed and WCL samples were resolved by imaged by immunoblotting against β-hexosaminidase subunit β. Data is representative of three independent experiments and quantification are shown in B. (C) The indicated cell lines were lysed and samples were split evenly between non-treated and PNGaseF treated. Asterisks denote deglycosylated protein (D). As described for panel C, except immunoblotting was against CI M6PR and the asterisks denote deglycosylated protein.

Supplemental Figure 4

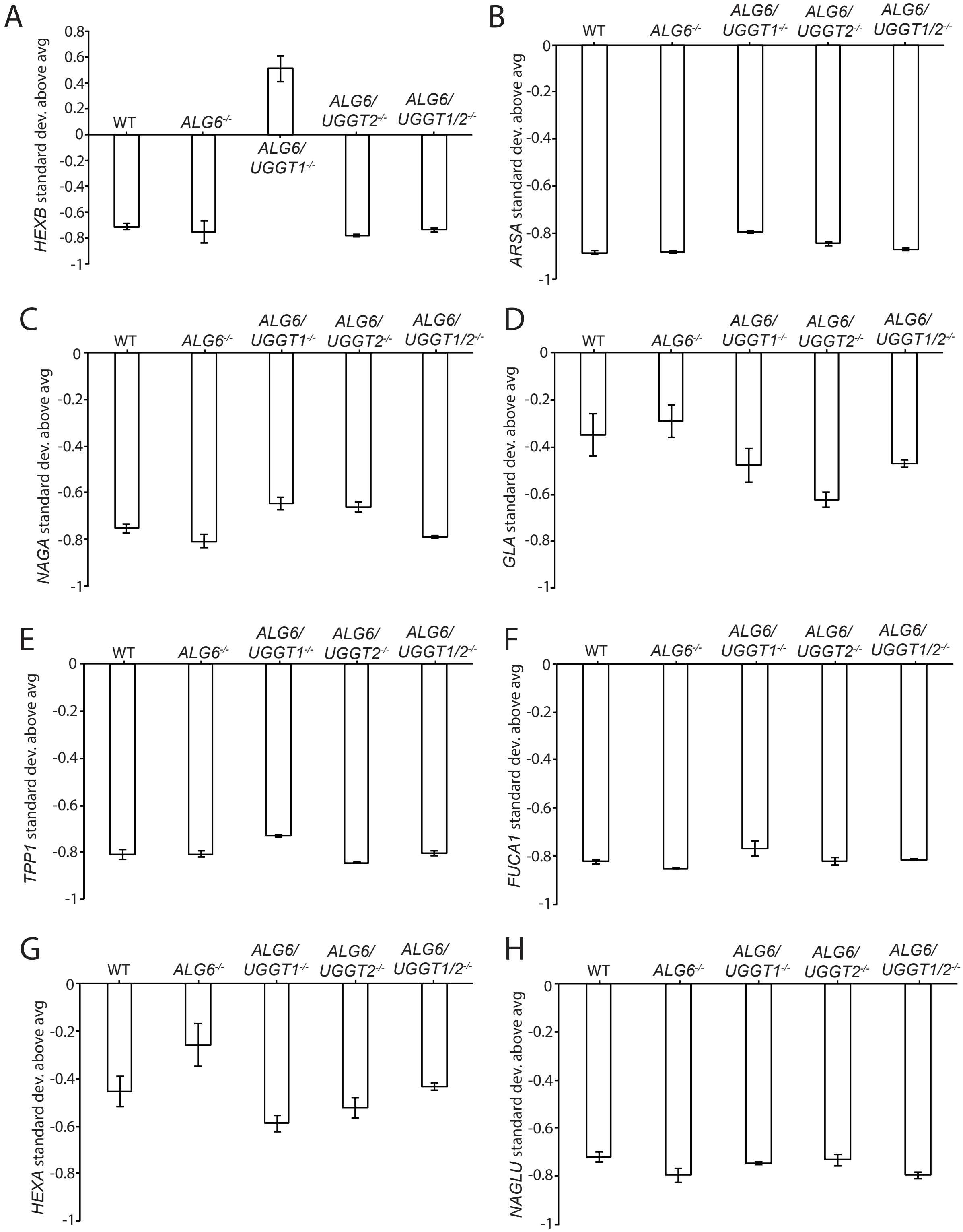


**Supplemental Figure 4. UPR induction in knock-out cell lines.**

(A) The indicated cells were lysed and WCL were resolved by SDS-PAGE and imaged by immunoblotting against BiP and GAPDH. DMSO was used as a vehicle control with tunicamycin (5 μg/ml) as a positive control. Data is representative of three independent experiments with quantification displayed in B. BiP expression levels were normalized to that of WT cells. And the corresponding GAPDH loading control. Error bars denote standard deviation. Asterisks denote a p-value of less than 0.05. (C) A subset of genes induced by the ATF6 UPR branch was analyzed by mRNA expression level in the denoted cell lines from RNAseq data presented in supplemental table 4. The standard deviation from the average expression level in counts per million in each cell line for all genes is plotted. Error bars represent standard deviation. Data are representative of three independent experiments. IRE1 (D) and PERK (E) induced genes characterized as described in C.

Supplemental Figure 5

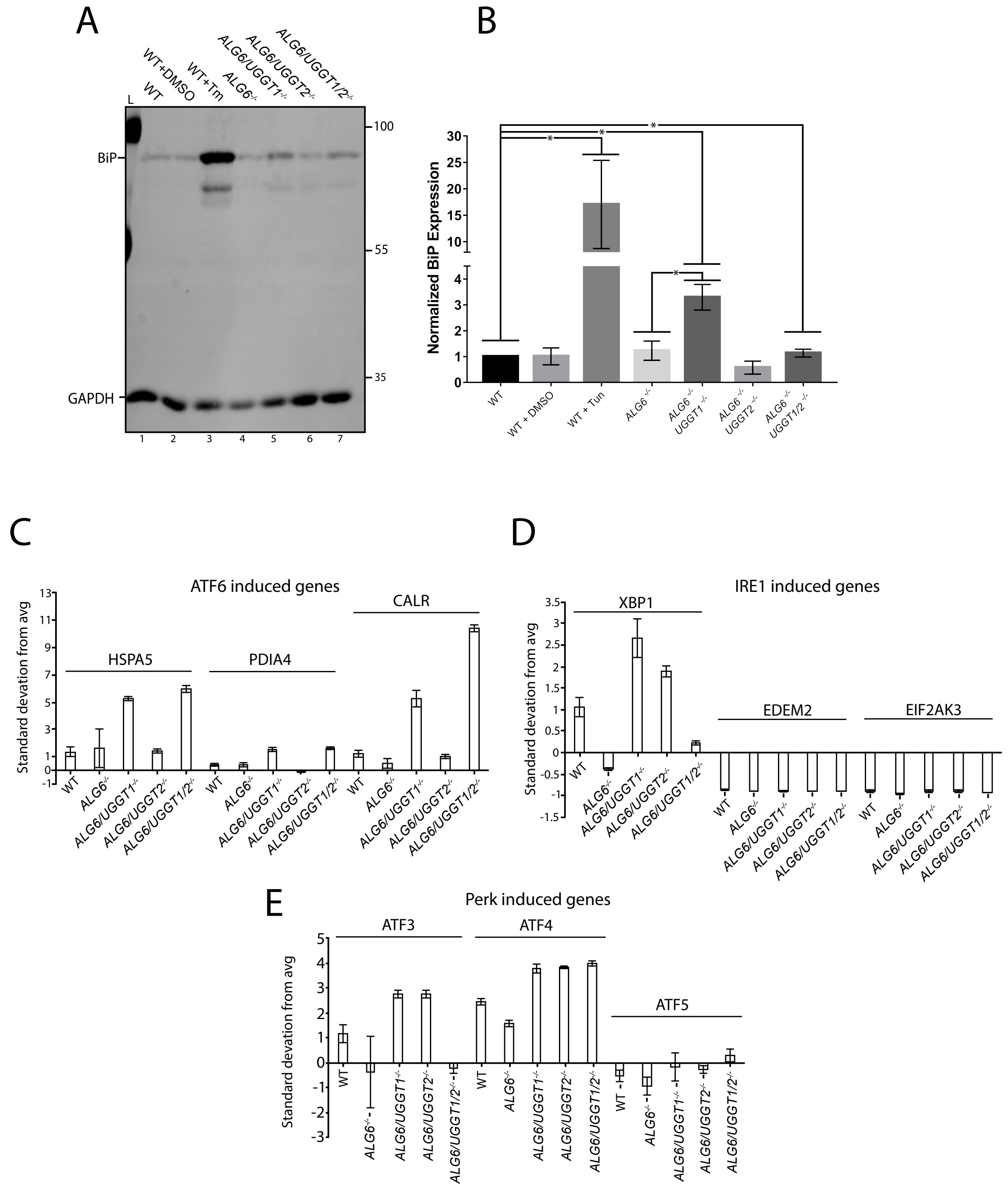


**Supplemental Figure 5. mRNA expression of lysosomal preferential UGGT2 substrates.**

(A) The expression of *HEXB* (A), *ARSA* (B), *NAGA* (C), *GLA* (D), *TPP1* (E), *FUCA1* (F), *HEXA* (G) and *NAGLU* (H) in the indicated cell lines was analyzed as described in Supplemental Figure 4C.
